# A PPARγ-like signalling axis integrates lipid homeostasis and mitochondrial quality control to preserve neuronal integrity

**DOI:** 10.64898/2026.09.28.755091

**Authors:** Dikaia Tsagkari, Maria Markaki, Tao Zhang, Ellen Nollen, Nektarios Tavernarakis

**Affiliations:** Division of Basic Sciences, School of Medicine, University of Crete, Heraklion 71003, Crete, Greece Figure; Institute of Molecular Biology and Biotechnology, Foundation for Research and Technology-Hellas, Heraklion 70013, Crete, Greece; European Research Institute for the Biology of Ageing, University Medical Centre Groningen, Antonius Deusinglaan 1, Groningen 9713 AV, The Netherlands; College of Food Science and Technology, Huazhong Agricultural University, Wuhan 430070, Hubei, China

## Abstract

Peroxisome proliferator-activated receptor gamma (PPARγ) is a central regulator of lipid metabolism and a critical mediator of neuroprotection, yet the mechanisms linking these functions remain unclear. Here, we identify the *Caenorhabditis elegans* nuclear receptor SEX-1 as a functional PPARγ-like regulator and define a pathway that coordinates lipid metabolism with mitochondrial homeostasis to preserve neuronal function. Integrating comparative genomics, multi-omics profiling, meta-analysis, and experimental validation, we show that SEX-1 orchestrates membrane sphingolipid metabolism, lipid droplet (LD) dynamics, and mitochondrial quality control through interconnected lipogenic and autophagic programs. Loss of *sex-1* disrupts this metabolic network, resulting in aberrant LD accumulation, mitochondrial dysfunction, and widespread alterations in sphingolipid metabolism. These defects compromise the structural integrity and thermosensory function of amphid neurons during early adulthood, while the capacity of SEX-1 to restrict neuronal lipid droplet accumulation declines with age. Notably, genetic modulation of sphingolipid metabolic pathways rescues neuronal defects, establishing a causal link between lipid metabolic homeostasis and neuronal maintenance. Together, our findings identify a PPARγ-like signalling axis that integrates sphingolipid metabolism, organelle quality control, and neuronal homeostasis, providing a mechanistic framework for understanding how metabolic regulation promotes neuroprotection.

## Introduction

Whole-body metabolism is coordinated by the peroxisome proliferator-activated receptors (PPARs), a superfamily of ligand-activated transcription factors that regulate lipid and glucose homeostasis^1–3^. Mammals express three PPAR isoforms with distinct physiological functions. PPARα primarily regulates fatty acid transport and mitochondrial β-oxidation, PPARγ controls adipogenesis, lipolysis and lipid storage, whereas PPARβ/δ coordinates both lipid and glucose metabolism^1, 2^. Despite their functional divergence, all three isoforms share an evolutionarily conserved modular architecture comprising an N-terminal ligand-independent transactivation domain (AF1), a highly conserved DNA-binding domain, and a C-terminal ligand-binding domain containing the ligand-dependent transactivation domain (AF2), which is required for nuclear localisation and protein-protein interactions^4–6^. Endogenous PPAR ligands include unsaturated FAs and lipid derivatives generated from dietary sources or FA catabolism. Ligand binding promotes transcriptional activation of target genes, while PPARs also repress gene expression through interactions with other transcription factors and regulatory proteins^1^.

Among the PPAR family, PPARγ has emerged as a central regulator of lipid and carbohydrate metabolism, with additional anti-inflammatory and neuroprotective functions in the nervous system^6, 7^. A major function of PPARγ is to promote the sequestration of FAs into lipid droplets (LDs) by regulating genes involved in lipid metabolism^8^. LDs are highly dynamic organelles that control lipid storage, synthesis, hydrolysis and trafficking^3, 9, 10^, and establish extensive contacts with the endoplasmic reticulum (ER), mitochondria, peroxisomes, Golgi apparatus and lysosomes^11^. In particular, dynamic interactions between LDs and mitochondria are essential for maintaining cellular energy homeostasis^10, 12^. During lipolysis or lipophagy, LD-derived FAs fuel mitochondrial β-oxidation and stimulate mitochondrial biogenesis, whereas mitochondria reciprocally regulate LD biogenesis and expansion^11, 13^. Although several candidate tethering proteins have been identified through interactomic studies, the molecular mechanisms governing LD-mitochondria interactions remain poorly understood^10, 11, 14^. Recent studies identified the LD-associated protein perilipin 5 (PLIN5) as a key regulator of mitochondrial biogenesis and PPAR signalling^15, 16^, suggesting reciprocal regulation between LD-mitochondria contacts and PPAR activity.

Pharmacological activation of PPARγ with thiazolidinediones (TZDs) alleviates neurodegeneration and promotes neuronal development in cultured neurons and mouse models of neurodegenerative disease^6, 17, 18^. However, the molecular mechanisms underlying PPARγ-mediated neuroprotection remain incompletely understood. To address this question, we sought to identify and characterise a functional PPARγ orthologue in *Caenorhabditis elegans*. The *C. elegans* genome encodes 284 nuclear hormone receptors (NHRs), most of which regulate development and lipid metabolism^19, 20^. Among these, NHR-49 has been established as the functional orthologue of PPARα, regulating multiple aspects of lipid metabolism, mitochondrial dynamics and lifespan, and mediating neuroprotection in models of Alzheimer’s disease^21, 22^. In contrast, a functional equivalent of PPARγ has not been identified.

Here, we identify the nuclear hormone receptor SEX-1, a previously characterised X-chromosome signal element required for sex determination^19, 23, 24^, as a functional orthologue of PPARγ in *C. elegans*. Integrating transcriptomic and lipidomic analyses, we demonstrate that SEX-1 regulates lipid metabolism by controlling the expression of genes involved in lipid biosynthesis and remodelling, thereby influencing the abundance of key lipid species, including sphingolipids. SEX-1 also regulates LD abundance across multiple tissues through SBP-1/SREBP1-dependent lipogenesis and lipophagy. Beyond lipid metabolism, SEX-1 coordinates mitophagy and complementary mitochondrial quality-control pathways, through the mitophagy receptor DCT-1/NIX, to maintain mitochondrial mass and function. This dual role of SEX-1 is linked to altered LD-mitochondria interactions involving the outer mitochondrial membrane translocase TOMM-20 and LD-associated proteins. Finally, we show that SEX-1 is required to preserve the integrity and function of the principal thermosensory neurons by regulating neuronal LD accumulation and the expression of the sphingolipid metabolism genes *mlt-8* and *sms-5*. Together, our findings establish SEX-1 as the functional PPARγ orthologue in *C. elegans* and reveal the molecular mechanisms through which PPARγ coordinates lipid homeostasis, mitochondrial quality control and neuroprotection.

## Results

### SEX-1 is a functional orthologue of PPARγ in *C. elegans*

To identify the *C. elegans* orthologue of human PPARγ, we performed BLASTp analysis using the human PPARγ protein (NCBI ID: P37231.3) as the query against the *C. elegans* proteome. This identified the steroid hormone receptor family member SEX-1 (also known as CELE_F44A6.2, NHR-24, CNR-14; NCBI ID: NP_001024662.1) as the top hit (72% similarity, E-value = 5e-35) (**Supplementary Table 1**). Reverse BLASTp analysis of SEX-1 against the human proteome identified PPARγ (NCBI ID: BAA23354.1) as the eleventh hit, with high sequence similarity (64% similarity, E-value = 3e-34) (**Supplementary Table 2**). Multiple sequence alignment using CLUSTAL W demonstrated extensive conservation of identical or functionally similar amino acids. Notably, the DNA-binding domain, including the P-box motif, was highly conserved, whereas the ligand-binding domain showed partial conservation, supporting functional similarity between the two proteins (**Figure S1A**).

To further assess structural conservation, we predicted the SEX-1 structure using AlphaFold^25^, and aligned it with human PPARγ using the flexible structural alignment algorithm FATCAT, which accounts for conformational flexibility and domain rearrangements during evolution^26^. The proteins exhibited significant structural similarity (P = 6.73e-06; FATCAT score = 822.35), with 425 equivalent residues, an RMSD of 6.98 Å, and four twists indicative of localized structural flexibility (**Figure S1B**). Three-dimensional visualization further demonstrated highly similar overall protein folding (**Figure S1C**).

Consistent with its role as a nuclear hormone receptor, SEX-1 localizes to nuclei and is expressed from oogenesis through mid-embryogenesis, with peak expression during early gastrulation^27^. To investigate its function, we examined *sex-1(y263)* mutants, which carry a G→A transition in the splice acceptor site at the intron 4–exon 5 junction downstream of the DNA-binding domain. This mutation disrupts normal splicing, generates frameshifted transcripts, and reduces SEX-1 protein abundance to undetectable levels by anti–SEX-1 antibody staining. As previously reported, *sex-1(y263)* mutants display dumpy morphology and egg-laying defects^23, 24^ (**Figure S1D**).

To experimentally evaluate the functional equivalence of SEX-1 and PPARγ, wild-type animals and *sex-1(y263)* mutants were treated with rosiglitazone, a potent PPARγ agonist (EC₅₀ = 43 nM for the human PPARγ ligand-binding domain). Molecular docking using AutoDock Vina^28, 29^ predicted comparable binding affinities of rosiglitazone for SEX-1 and human PPARγ (**Movie S1**, **Movie S2**), with binding energies of −6.363 kcal/mol and −6.532 kcal/mol, respectively. We next identified established PPARγ target genes from publicly available ChIP-seq datasets^30^, and examined the expression of their *C. elegans* orthologues following rosiglitazone treatment. Whereas 0.1 μM and 1 μM rosiglitazone produced no detectable transcriptional changes, treatment with 10 μM significantly induced target gene expression in day 1 adults (**Figure S1E**). Specifically, rosiglitazone increased expression of *lgg-1* (Gabarap), *dgtr-1* (Dgat2), and *tomm-40* (Tomm40) in wild-type animals but not in *sex-1(y263)* mutants, demonstrating that these transcriptional responses require SEX-1 (**Figure 1A**).

**Figure 1.**
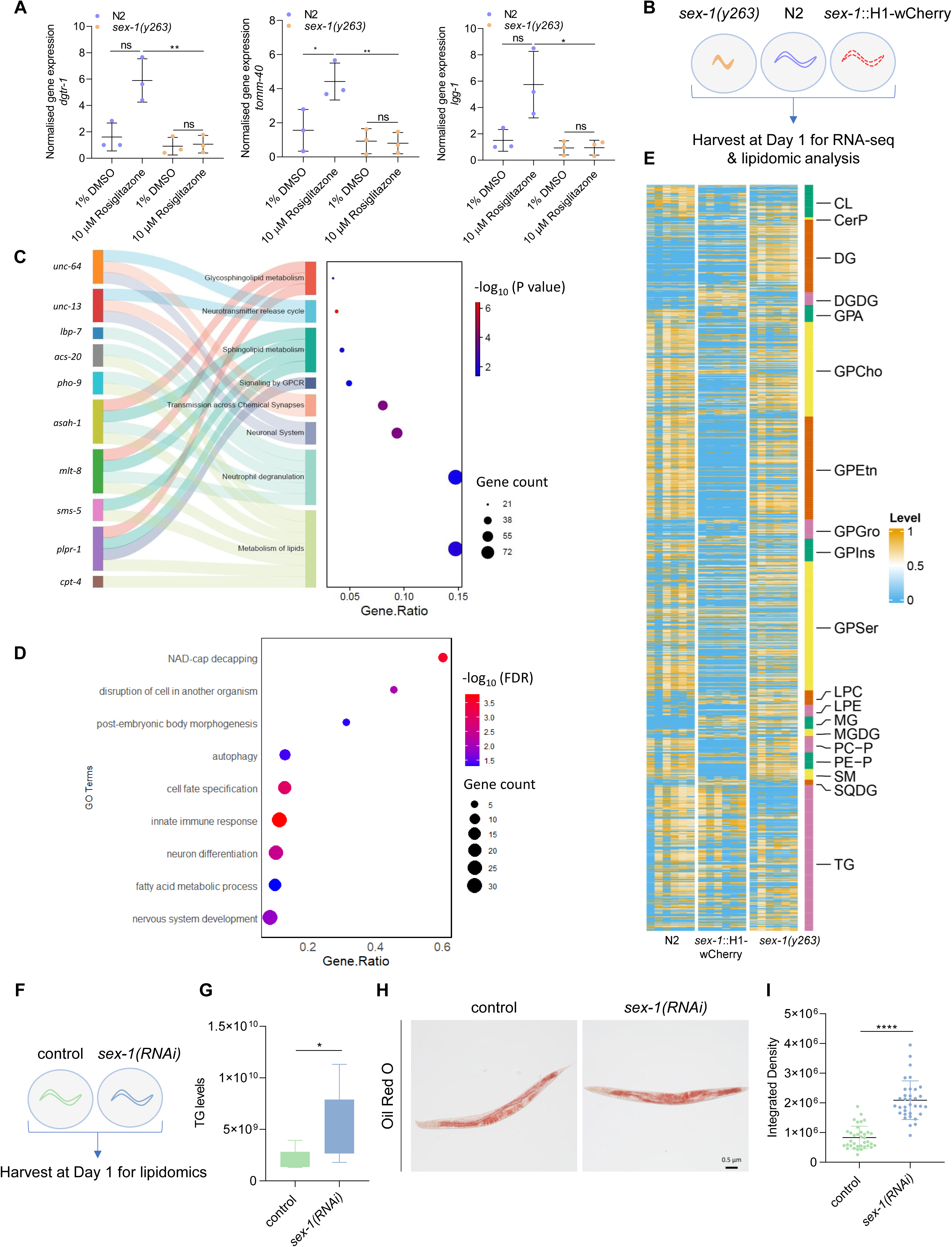
SEX-1 modulates lipid metabolism and the organismal lipid profile. A. Normalised expression levels of *dgtr-1*, *tomm-40* and *lgg-1* in wild-type (N2) animals and *sex-1(y263)* mutants treated with 1% DMSO or 10 µM Rosiglitazone (n=3 independent experiments, *P < 0.05, **P < 0.01; unpaired t-test). B. Experimental set-up for transcriptomic and lipidomic analysis. C. Sankey and dot plot of differentially expressed genes (DEGs) enriched in gene ontology (GO) analysis of *sex-1(y263)* mutants. The Sankey map on the left shows the genes associated with more than one GO term. The horizontal coordinate of the dot plot on the right is the gene ratio of each GO team, the colour of each circle represents the -log_10_ (P value) and the size of each circle represents the number of genes enriched in each GO term. D. Dot plot of enriched terms in gene ontology (GO) analysis from differentially expressed genes (DEGs) of *sex-1*::H1-wCherry transgenic animals. The horizontal coordinate of the dot plot is the gene ratio of each GO term, the colour of each circle represents the -log_10_ (FDR) and the size of each circle represents the number of genes enriched in each GO term. E. Heatmap of lipidomic analysis in *sex-1(y263)* mutants, wild-type (N2) and *sex-1*::H1-wCherry animals, showing differential abundance across all lipid classes. Colour intensity represents the relative changes in lipid levels, with orange indicating upregulation and blue indicating downregulation. F. Experimental set-up for follow-up lipidomic analysis. G. Boxplot showing changes in triglyceride (TG) levels in wild-type (N2) animals treated with control or *sex-1* RNAi (n=6 technical replicates per group, 3000 worms per replicate, *P < 0.05; two-tailed unpaired t-test). H. Representative images of wild-type (N2) animals treated with control or *sex-1* RNAi and stained with Oil Red O (Images were acquired using a ×4 objective lens, n=3 independent experiments). I. Quantification of Oil Red O staining as shown in (H) (n=3 independent experiments, 90 animals in total, ****P < 0.0001; two-tailed unpaired t-test).

To further investigate ligand recognition, we used AlphaFold 3^31^ to model the interaction between SEX-1 and oleic acid, an endogenous unsaturated fatty acid and natural ligand of PPARγ (**Movie S3**). The predicted complex yielded a template modelling (pTM) score of 0.53, indicating a plausible overall fold, and an interface pTM (ipTM) score of 0.83, supporting high confidence in the predicted protein–ligand interaction.

To determine whether SEX-1 regulates biological processes similar to mammalian PPARγ, we performed genome-wide transcriptomic analysis (**Figure 1B**) and compared the *sex-1(y263)* transcriptome with published datasets from PPARγ-deficient multipotent mesenchymal stromal cells^32^. This analysis identified 748 shared Gene Ontology (GO) terms (**Figure S1F**), indicating substantial conservation of downstream biological pathways. The most significantly enriched shared GO categories included growth and development, neuronal signalling, metabolism, muscle development, autophagy, and stress responses (**Figure S1G**).

To place SEX-1 in an evolutionary context, we generated a maximum-likelihood phylogenetic tree including the three human PPAR isoforms (PPARα, PPARδ, and PPARγ), *C. elegans* nuclear hormone receptors identified as candidate PPARγ orthologues by BLASTp, and NHR-49, the established functional orthologue of PPARα. Phylogenetic analysis showed that SEX-1 clustered closest to the human PPAR clade, indicating greater evolutionary similarity to vertebrate PPARs than the other *C. elegans* nuclear hormone receptors examined (NHR-23, NHR-35, NHR-49, and NHR-85) (**Figure S1H**). Together, these structural, phylogenetic, and functional analyses identify SEX-1 as the strongest candidate functional orthologue of mammalian PPARγ in *C. elegans*.

To determine whether SEX-1 and NHR-49/PPARα regulate overlapping transcriptional programs, we performed a meta-analysis comparing differentially expressed genes (DEGs) (**Supplementary Table 3**) from the *sex-1(y263)* transcriptome with published DEGs (log₂ fold change ≥ 1 or ≤ −1) from *nhr-49(nr2041)* mutants^33^. The two datasets shared only 8.4% of DEGs (**Figure S2A**), indicating that SEX-1 and NHR-49 regulate largely distinct transcriptional programs. This parallels the functional relationship between mammalian PPARγ and PPARα^34^, while suggesting a limited set of common regulatory targets. GO enrichment analysis of the shared genes revealed significant enrichment for pathways involved in environmental adaptation, development, and neurogenesis, consistent with a role in coordinating metabolic and stress responses (**Figure S2B**).

Collectively, these bioinformatic, structural, phylogenetic, and functional analyses identify SEX-1 as a structural and functional orthologue of mammalian PPARγ in *C. elegans*.

### SEX-1 orchestrates neuronal gene expression and systemic lipid metabolism

Having identified SEX-1 as a functional PPARγ orthologue, we next investigated its role in transcriptional regulation and metabolic homeostasis. To this end, we further analysed our genome-wide transcriptomic dataset after confirming that *sex-1* expression was undetectable in *sex-1(y263)* mutants following 40 PCR cycles, whereas *sex-1* mRNA levels were elevated in *sex-1*::H1-wCherry transgenic animals (**Figure S2C**). Whole-genome sequencing of the transgenic strain identified a predominant integration event on the X chromosome, upstream of the first exon of *sex-1* (**Figure S2D**), providing a molecular explanation for the increased *sex-1* expression. Unsupervised principal component analysis (PCA) demonstrated clear separation between wild-type animals and age-matched *sex-1(y263)* mutants (**Figure S2E**). Differential expression analysis identified 3,125 upregulated and 370 downregulated genes in 1-day-old adult mutants, of which 1,417 and 41 genes, respectively, exhibited changes of at least twofold (**Figure S2F**) (**Supplementary Table 3**). Gene Ontology (GO) enrichment analysis revealed significant overrepresentation of pathways associated with neuronal signalling and lipid metabolism (**Figure 1C**).

Similarly, PCA distinguished *sex-1*::H1-wCherry transgenic animals from wild-type controls (**Figure S2G**). Compared with wild-type, transgenic animals displayed 378 upregulated and 85 downregulated genes, including 117 and 19 genes, respectively, with changes ≥2-fold (**Figure S2H**) (**Supplementary Table 4**). Enrichment analysis highlighted pathways involved in neurodevelopment and stress responses associated with neuroprotective adaptation (**Figure 1D**). Collectively, these transcriptomic analyses demonstrate that SEX-1 coordinates transcriptional programmes governing neuronal function and lipid metabolism, revealing broader regulatory roles beyond sex determination that are consistent with PPARγ signalling networks^3, 35^.

To determine whether these transcriptional changes were accompanied by metabolic remodelling, we performed lipidomic profiling of 1-day-old adult wild-type animals, *sex-1*::H1-wCherry transgenics and *sex-1(y263)* mutants (**Figure 1B**). PCA revealed distinct genotype-specific lipidomic signatures (**Figure S3A**). Quantitative analysis of lipid species normalised to total protein content identified significant alterations across multiple lipid classes (**Figure 1E**). *sex-1(y263)* mutants exhibited markedly reduced triacylglycerol (TG) levels accompanied by increased diacylglycerol (DG) and monoacylglycerol (MG) levels (**Figure S3B**). In contrast, *sex-1*::H1-wCherry animals showed no significant changes in TG or DG abundance but displayed an overall lipid profile opposite to that of *sex-1(y263)* mutants, with the exception of MGs, which were elevated in both genotypes (**Figure S3B**). This pattern is consistent with altered lipolytic activity, whereby TGs are hydrolysed sequentially to DGs and MGs through neutral lipid droplet-associated or lysosomal (acid) lipolysis, releasing fatty acids and glycerol^36^.

Beyond neutral lipids, *sex-1(y263)* mutants displayed significant increases in two phospholipid classes together with sphingomyelin, the predominant sphingolipid species (**Figure S3C**). Although phospholipid abundance was not significantly altered in transgenic animals relative to wild-type, their levels were consistently lower than those observed in *sex-1(y263)* mutants, indicating an opposing lipidomic phenotype (**Figure S3C**). Furthermore, all glycerophospholipid subclasses, major structural membrane lipids that are particularly abundant in the nervous system and essential for cellular compartmentalisation^37^, were significantly reduced in transgenic animals relative to wild-type (**Figure S3D**). Direct comparison of glycerophosphoethanolamine (GPE), glycerophosphoglycerate (GPG) and diacyl glycerophosphatidic acid (GPA) between *sex-1(y263)* mutants and transgenic animals further emphasised their contrasting lipid profiles (**Figure S3D**), in agreement with GO enrichment implicating glycerophospholipid metabolism. Given that phospholipids and sphingolipids function as key signalling molecules in neuronal tissues^38^, these findings support a role for SEX-1 in regulating neuronal lipid signalling, consistent with the transcriptomic data.

Because *sex-1(y263)* mutants exhibit severe embryonic defects and reduced viability, we next examined animals subjected to post-embryonic *sex-1* knockdown by RNA interference (RNAi; **Figure S3E**). Lipidomic analysis of 1-day-old adult wild-type and *sex-1(RNAi)* animals (**Figure 1F**) revealed distinct clustering of the two conditions (**Figure S3F**). However, no significant differences were detected across most lipid classes, with the exception of a modest but significant increase in TG abundance following *sex-1* knockdown (**Figure 1G**; **Figure S3G**). Oil Red O staining (ORO; indicator of fat storage and overall triglyceride levels^39^) confirmed increased fat accumulation in *sex-1(RNAi)* animals (**Figure 1H,I**), and direct biochemical quantification of TGs normalised to protein content independently validated this increase (**Figure S3H**).

We also performed ORO staining on the genotypes included in the multi-omics analyses (**Figure S3I**). Consistent with the lipidomic data, *sex-1(y263)* mutants exhibited almost undetectable TG stores, whereas *sex-1*::H1-wCherry animals displayed reduced fat accumulation relative to wild-type (**Figure S3J**). The contrasting TG phenotypes observed between *sex-1(y263)* mutants and *sex-1(RNAi)* animals likely reflect the developmental expression pattern of *sex-1*, which is highest during embryogenesis and declines during larval and adult stages, whereas RNAi-mediated silencing was initiated post-embryonically. Supporting this interpretation, RNAi treatment over two consecutive generations recapitulated the lipid phenotype of *sex-1(y263)* mutants (**Figure S3K**).

This developmental model is further supported by transcriptomic analysis showing reduced expression of *vit-1* and *vit-3* in *sex-1(y263)* mutants. These genes encode vitellogenin yolk proteins that assemble into high-density lipoprotein particles serving as major lipid storage structures in *C. elegans*^40^. Loss of vitellogenins promotes intestinal atrophy by driving the mobilisation of maternal lipid reserves to developing progeny^41, 42^.

Together, these findings establish SEX-1 as a central regulator of systemic lipid homeostasis that integrates developmental timing with transcriptional programmes controlling neuronal function and lipid metabolism.

### SEX-1 regulates lipid droplet abundance through SBP-1-dependent lipogenesis and enhanced lipophagy

Lipid droplets (LDs) are dynamic organelles that store triglycerides and vary in size and metabolic activity. Their surface is composed of a phospholipid monolayer decorated with a distinct repertoire of proteins, including perilipins (PLINs)^43^. To investigate whether SEX-1 regulates LD dynamics, we performed super-resolution confocal microscopy in transgenic animals ubiquitously expressing PLIN-1 (formerly MDT-28)^44^, following *sex-1 RNAi* (**Figure 2A**). Quantitative analysis revealed a significant increase in both LD number and size in *sex-1(RNAi)* animals compared with controls (**Figure 2B**).

**Figure 2.**
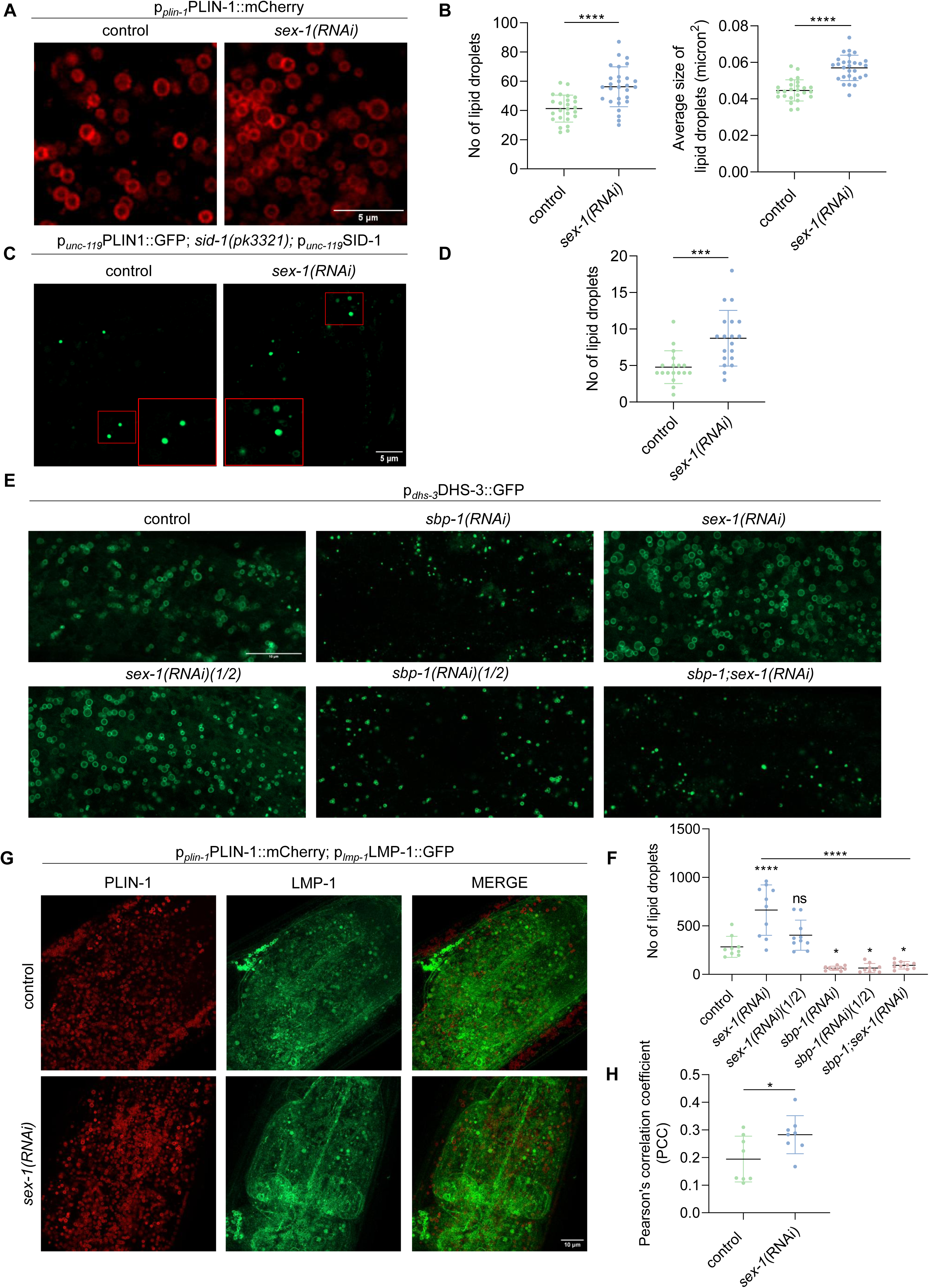
SEX-1 regulates lipid droplet abundance via SBP-1-regulated lipogenesis and lipophagy. A. Representative zoomed-in images of lipid droplets in the head of transgenic p*_plin-1_*PLIN-1::mCherry animals treated with control or *sex-1* RNAi (Images were acquired using a ×63 objective lens, n=3 independent experiments). B. Number and average size of lipid droplets in a 15 x 15 µm area in the head of animals assessed by fluorescence as shown in (A) following treatment with control or *sex-1* RNAi (n= 3 independent experiments, 25 animals in total, ****P < 0.0001; two-tailed unpaired t-test). Each dot represents the number or average size of lipid droplets in a specific area of an individual worm. C. Representative images of lipid droplets in the nerve ring of transgenic p*_unc-119_*PLIN1::GFP animals treated with control or *sex-1* RNAi (Images were acquired using a ×63 objective lens, n=3 independent experiments). D. Number of lipid droplets in the nerve ring of animals assessed by fluorescence as shown in (C) following treatment with control or *sex-1* RNAi (n=3 independent experiments, 23 animals in total, ***P < 0.001; two-tailed unpaired t-test). Each dot represents the number of lipid droplets in the nerve ring of an individual worm. E. Representative images of lipid droplets in a 50 x 25 µm area in the intestine of transgenic p*_dhs-3_*DHS-3::GFP animals treated with control, *sbp-1*, *sex-1*, *sex-1(1/2)*, *sbp-1(1/2)* or *sbp-1;sex-1* RNAi (Images were acquired using a ×63 objective lens, n=1 experiment). F. Number of intestinal lipid droplets assessed by fluorescence as shown in (E) following treatment with control, *sbp-1*, *sex-1*, *sex-1(1/2)*, *sbp-1(1/2)* or *sbp-1;sex-1* RNAi (n=1 experiment, 10 animals, *P < 0.05, ****P < 0.0001; one-way analysis of variance (ANOVA)). Each dot represents the number of lipid droplets in the intestine of an individual worm. G. Representative images of transgenic animals that co-express PLIN-1 (mCherry) and LMP-1 (GFP) showing their localisation after treatment with control or *sex-1* RNAi (Images were acquired using a ×63 objective lens, n=2 independent experiments). H. Pearson’s correlation coefficient values after measuring the colocalization between PLIN-1 and LMP-1 (n=2 independent experiments, 20 animals in total, *P < 0.05; two-tailed unpaired t-test).

Given that our transcriptomic and lipidomic analyses implicated SEX-1 in neuronal lipid metabolism, we next examined neuronal LD dynamics using RNAi-sensitive *sid-1(pk3321); punc-119*::SID-1 animals expressing a pan-neuronal GFP fusion of *Drosophila* PLIN1^45^ (**Figure 2C**). Super-resolution imaging demonstrated a significant increase in neuronal LD abundance following *sex-1* knockdown (**Figure 2D**), indicating that SEX-1 regulates LD homeostasis across multiple tissues, including neurons.

To further assess the role of SEX-1 in intestinal LD regulation, we analysed transgenic animals expressing the LD-associated protein dehydrogenase-3 (DHS-3) fused to GFP^44^ (**Figure S4A**). Consistent with our previous observations, *sex-1 RNAi* significantly increased LD abundance, as quantified by mean DHS-3::GFP fluorescence intensity (**Figure S4B**). Conversely, transgenic *sex-1*::H1-wCherry animals expressing DHS-3::GFP exhibited reduced LD abundance relative to controls (**Figure S4A**, **B**). Notably, RNAi-mediated depletion of *sex-1* in these double-transgenic animals restored LD abundance to control levels (**Figure S4A**, **B**), providing further evidence that SEX-1 negatively regulates LD accumulation.

To investigate the molecular mechanisms underlying the increase in LD abundance following *sex-1* depletion, we examined the expression of genes involved in lipogenesis, lipophagy, and lipolysis, including candidates identified in the transcriptomic analysis of *sex-1(y263)* mutants. In 1-day-old adult *sex-1(RNAi)* animals, expression of the lipogenic regulator *sbp-1*, the stearoyl-CoA desaturase genes *fat-6* and *fat-7*, and the lysosomal lipases *lipl-1* and *lipl-3* was significantly upregulated, whereas expression of genes associated with lipolysis remained unchanged (**Figure S4C**).

Because SBP-1 is the *C. elegans* orthologue of mammalian SREBP1 (sterol regulatory element-binding protein 1) and is required for transcriptional activation of *fat-6* and *fat-7*^46^, we next assessed its contribution to SEX-1-dependent LD regulation. RNAi-mediated knockdown of *sbp-1* reduced LD abundance in *pdhs-3*::DHS-3::GFP animals (**Figure S4D**) and partially suppressed the LD accumulation induced by *sex-1* RNAi (**Figure S4E**). These findings were confirmed by super-resolution confocal microscopy (**Figure 2E**, **F**; **Figure S4F**), demonstrating that SBP-1 is required for the lipogenic response associated with *sex-1* depletion.

To determine whether altered LD turnover also contributes to this phenotype, we assessed lipophagy using transgenic animals co-expressing PLIN-1::mCherry and the lysosomal marker LMP-1::GFP (**Figure 2G**). Colocalization between LDs and lysosomes, quantified using Pearson’s correlation coefficient (PCC), was significantly increased following *sex-1* knockdown (**Figure 2H**), indicating enhanced lipophagic flux.

Collectively, these findings demonstrate that loss of SEX-1 increases LD abundance through SBP-1-dependent activation of lipogenesis, while simultaneously enhancing lipophagy, suggesting that SEX-1 coordinates lipid storage and turnover to maintain LD homeostasis.

### SEX-1 regulates mitochondrial homeostasis through DCT-1-dependent mitophagy and complementary mitochondrial quality control pathways

PPARγ is a central regulator of energy metabolism and has been implicated in the transcriptional control of numerous mitochondrial genes^5^. Consistent with this role, our transcriptomic analysis identified differential expression of multiple mitochondria-associated genes following modulation of SEX-1. To investigate whether SEX-1 regulates mitochondrial homeostasis, we first examined the effects of *sex-1* downregulation and overexpression on mitochondrial abundance in different tissues (**Figure 3A**, **Figure S5A**). In both the intestine and body wall muscle, mitochondrial mass was significantly increased in *sex-1(RNAi)* animals compared with controls, whereas it was reduced in *sex-1*::H1-wCherry transgenic animals overexpressing *sex-1*, indicating that SEX-1 negatively regulates mitochondrial abundance (**Figure 3B**, **Figure S5B**). Furthermore, *sex-1* knockdown rescued the reduction in mitochondrial mass observed in *sex-1*::H1-wCherry animals, further supporting a direct role for SEX-1 in regulating mitochondrial content (**Figure 3B**, **Figure S5B**).

**Figure 3.**
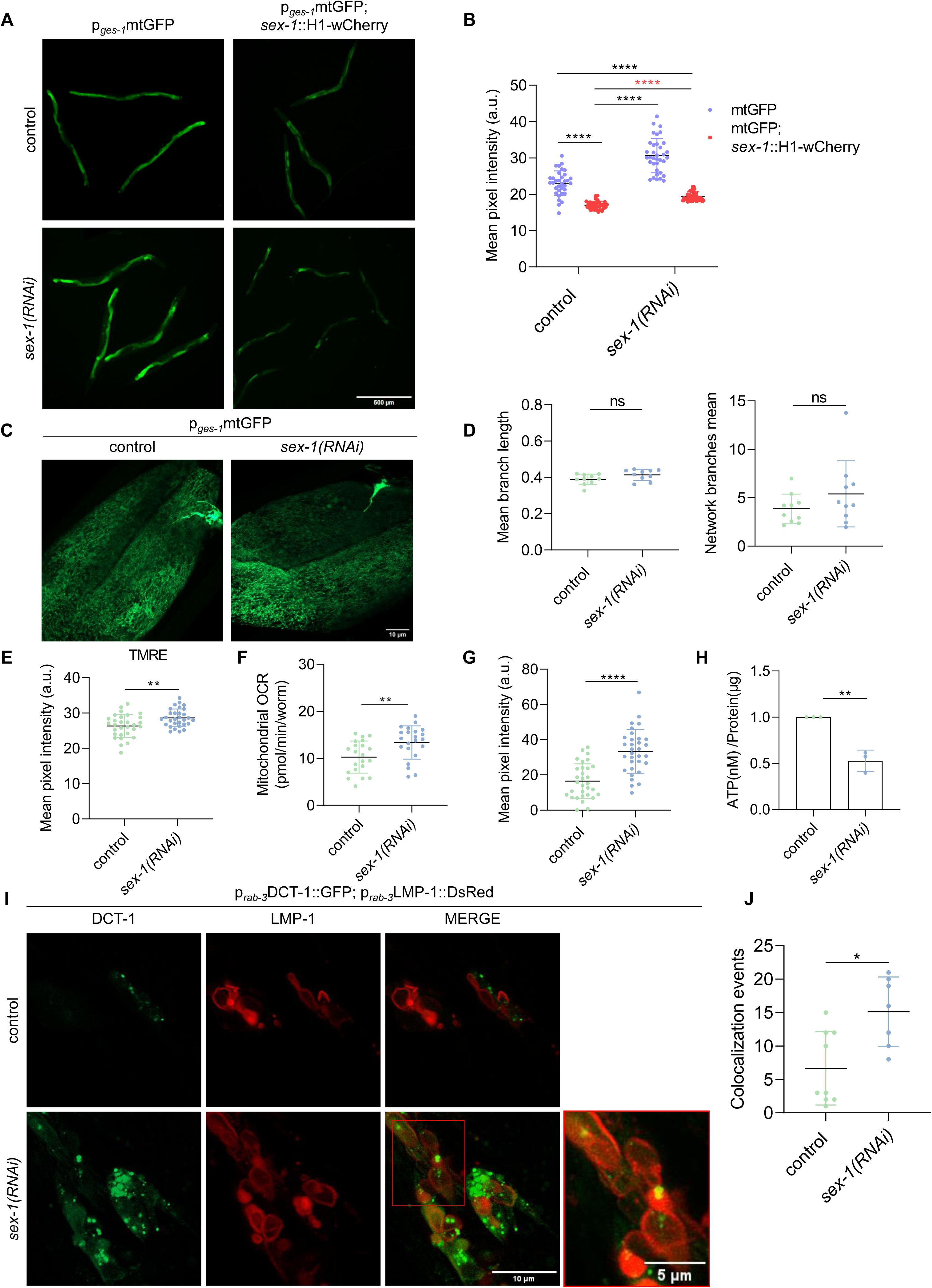
SEX-1 modulates mitochondrial abundance and function by coordinating DCT-1-dependent mitophagy. A. Representative images of transgenic *sex-1*::H1-wCherry and/or p*_ges-1_*mtGFP animals treated with control or *sex-1* RNAi showing intestinal mitochondrial mass (Images were acquired using a ×4 objective lens, n=3 independent experiments). B. Quantification of the mean GFP pixel intensity in the intestinal mitochondrial matrix of animals as shown in (A) (n=3 independent experiments, 90 animals in total, ****P < 0.0001; one-way analysis of variance (ANOVA)). C. Representative images of transgenic p*_ges-1_*mtGFP animals treated with control or *sex-1* RNAi showing mitochondrial morphology in the intestine (Images were acquired using a ×63 objective lens, n=3 independent experiments). D. Quantification of the mean branch length and mitochondrial network branches in transgenic p*_ges-1_*mtGFP animals as shown in (C) (n=3 independent experiments, 30 animals in total, two-tailed unpaired t-test). E. Quantification of the mean pixel signal of TMRE staining showing the mitochondrial membrane potential in wild-type (N2) animals treated with control or *sex-1* RNAi (n=3 independent experiments, 90 animals in total, **P < 0.01; two-tailed unpaired t-test). F. Mitochondrial oxygen consumption rates of wild-type (N2) animals treated with control or *sex-1* RNAi (n=3 independent experiments, 60 animals in total, **P < 0.01; two-tailed unpaired t-test). G. Mitochondrial ROS levels in wild-type (N2) animals treated with control or *sex-1* RNAi (n=3 independent experiments, 90 animals in total, ****P < 0.0001; two-tailed unpaired t-test). H. Quantification of ATP levels in whole wild-type (N2) animals treated with control or *sex-1* RNAi (n=3 independent biological replicates examined in one experiment, **P < 0.01; two-tailed unpaired t-test). I. Representative images of transgenic animals co-expressing DCT-1 (GFP) and LMP-1 (DsRed) in neurons showing their localisation after treatment with control or *sex-1* RNAi (Images were acquired using a ×63 objective lens, n=3 independent experiments). J. Quantification of colocalization events between DCT-1 and LMP-1 in transgenic animals treated with control or *sex-1* RNAi. Increased colocalization indicates mitophagy stimulation (n=3 independent experiments, 25 animals in total, *P < 0.05; two-tailed unpaired t-test).

We next assessed mitochondrial network integrity and found that mitochondrial morphology and network architecture were preserved in *sex-1(RNAi)* animals across tissues compared with age-matched controls (**Figure 3C**, **D**; **Figure S5C**, **D**). To determine whether these changes were accompanied by altered mitochondrial function, we measured mitochondrial membrane potential using the fluorescent dye TMRE (tetramethylrhodamine ethyl ester perchlorate) (**Figure S5E**). SEX-1 depletion increased mitochondrial membrane potential (**Figure 3E**) and oxygen consumption (**Figure 3F**). In parallel, MitoTracker™ Red CMXRos staining revealed elevated mitochondrial reactive oxygen species (ROS) production (**Figure 3G**, **Figure S5F**), while intracellular adenosine triphosphate (ATP) levels were reduced (**Figure 3H**). The combination of increased oxygen consumption, elevated ROS production, and reduced ATP levels is indicative of impaired mitochondrial efficiency and dysfunction^47^. Collectively, these findings demonstrate that SEX-1 regulates both mitochondrial abundance and functional homeostasis.

Transcriptomic analysis further revealed that expression of the mitophagy receptor DCT-1/NIX^48^ was upregulated in *sex-1(y263)* mutants. To validate this observation, we monitored DCT-1 expression using a DCT-1::GFP reporter following *sex-1* RNAi. Consistent with the transcriptomic data, DCT-1::GFP fluorescence was significantly increased in SEX-1-depleted animals relative to controls (**Figure S5G**, **5H**). We next assessed mitophagic activity *in vivo* using the mitochondria-targeted Rosella (mtRosella) biosensor (**Figure S5I**). SEX-1 depletion significantly decreased the GFP/DsRed fluorescence ratio, consistent with enhanced mitophagy (**Figure S5J**). To confirm activation of DCT-1-dependent mitophagy, we performed confocal microscopy in transgenic animals co-expressing DCT-1::GFP and the lysosomal marker LMP-1::DsRed following *sex-1* RNAi (**Figure 3I**). Quantification of DCT-1/LMP-1 colocalization revealed a significant increase in mitophagic events in neurons of *sex-1(RNAi)* animals compared with controls (**Figure 3J**).

Notably, enhanced mitophagy occurred despite the increased mitochondrial mass observed following SEX-1 depletion, indicating that SEX-1 regulates multiple, and potentially opposing, aspects of mitochondrial homeostasis. To identify additional pathways contributing to this phenotype, we examined the expression of other mitochondrial quality control regulators and found that *sir-2.2*, the *C. elegans* orthologue of mammalian SIRT4, was downregulated following *sex-1* RNAi (**Figure S6A**). Previous studies have shown that SIRT4 deficiency promotes fatty acid oxidation and enhances mitochondrial gene expression in mouse liver and muscle cells^49^. Consistent with these findings, *sir- 2.2* knockdown increased mitochondrial content in p*_ges-1_*mtGFP animals, whereas simultaneous silencing of *sir-2.2* and *sex-1* did not further increase mitochondrial abundance, suggesting that SEX-1 regulates mitochondrial content, at least in part, through *sir-2.2* (**Figure S6B**, **C**). Importantly, SEX-1 depletion did not affect the mtDNA/nDNA ratio (**Figure S6D**), indicating that the increase in mitochondrial abundance occurs independently of detectable changes in mitochondrial DNA copy number.

Together, these results identify SEX-1 as a regulator of mitochondrial homeostasis that coordinates DCT-1-dependent mitophagy with additional mitochondrial quality control pathways to modulate mitochondrial abundance and function.

### SEX-1 regulates mitochondria–lipid droplet contacts through TOMM-20

Physical contacts between lipid droplets (LDs) and mitochondria are critical for coordinating energy metabolism, lipid storage, and lipolysis^12, 16^. To investigate these contacts *in vivo*, we generated transgenic animals co-expressing PLIN-1::mCherry and mito::GFP. Super-resolution confocal microscopy revealed frequent LD–mitochondria contacts in both body wall muscle (BWM) and intestinal cells (**Figure 4A**, **B**). Quantitative distance analysis in BWM demonstrated that *sex-1* downregulation significantly increased LD–mitochondria contacts, resulting in more than a twofold increase in the number of mitochondria contacting LDs compared with controls (**Figure 4C**). Consistent with these findings, intestinal cells from *sex-1(RNAi)* animals exhibited 919 LD– mitochondria contacts compared with 474 in control animals (**Figure 4D**). These results demonstrate that SEX-1 regulates not only the abundance of LDs and mitochondria but also the frequency of their contacts.

**Figure 4.**
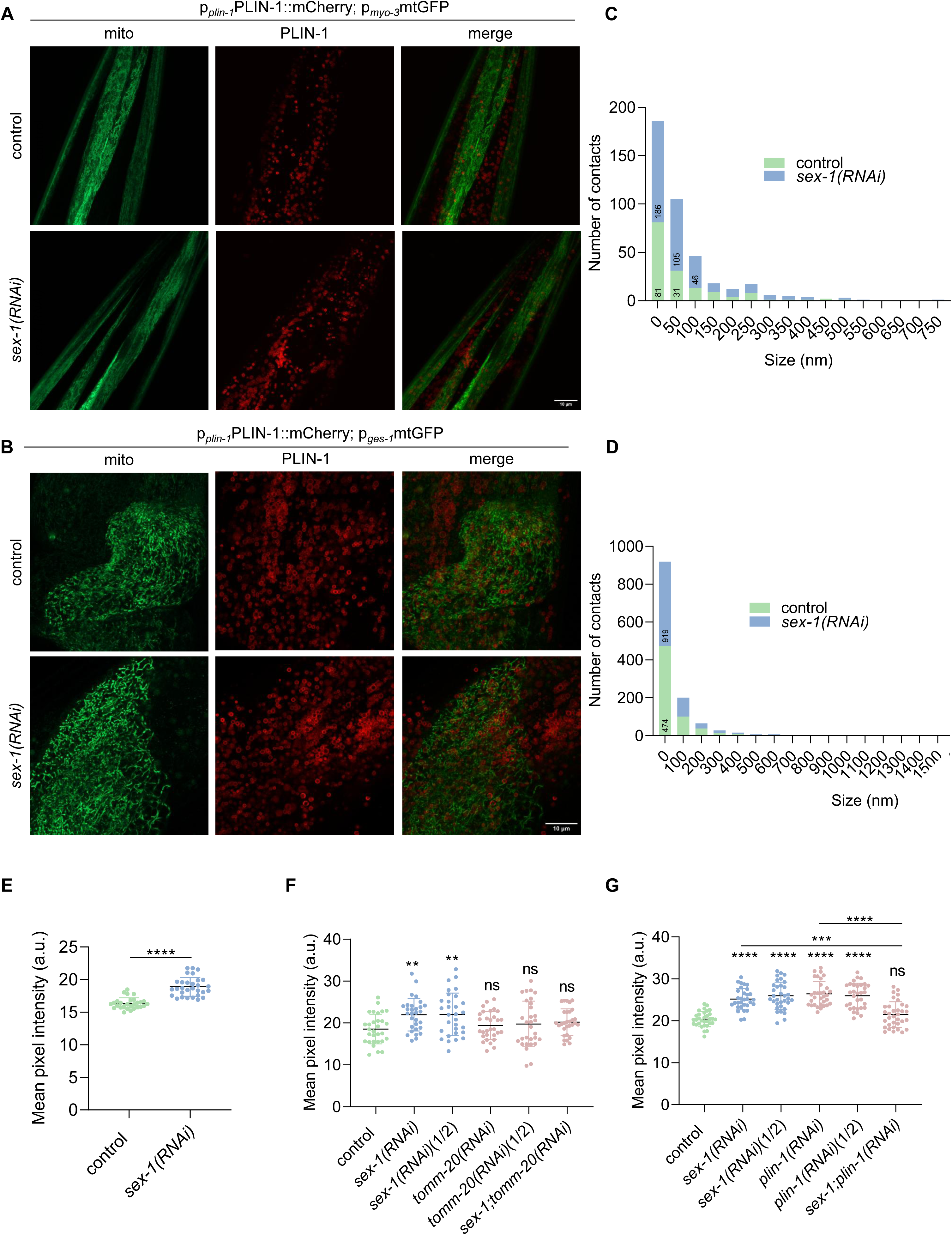
SEX-1 regulates mitochondria-lipid droplet interactions. A. Localisation of mitochondria (GFP) and lipid droplets (mCherry) in the body wall muscle (BWM) of transgenic animals treated with control or *sex-1* RNAi (Images were acquired using a ×63 objective lens, n=3 independent experiments). B. Localisation of mitochondria (GFP) and lipid droplets (mCherry) in the intestine of transgenic animals treated with control or *sex-1* RNAi (Images were acquired using a ×63 objective lens, n=3 independent experiments). C. Histogram showing the changes in contact length between mitochondria and lipid droplets in the BWM as determined in (A) (n=3 independent experiments, 30 animals in total). D. Histogram showing the changes in contact length between mitochondria and lipid droplets in the intestine as determined in (B) (n=3 independent experiments, 30 animals in total). E. Quantification of the mean mRFP fluorescence signal in transgenic p*_myo-3_*TOMM-20::mRFP animals treated with control or *sex-1* RNAi (n=3 independent experiments, 90 animals in total, ****P < 0.0001; Mann-Whitney test). F. Quantification of the mean GFP pixel signal in transgenic p*_dhs-3_*DHS-3::GFP animals treated with control, *tomm-20*, *sex-1*, *sex-1(1/2)*, *tomm-20(1/2)* or *tomm-20;sex-1* RNAi (n=3 independent experiments, 90 animals in total, **P < 0.01; one-way analysis of variance (ANOVA)). G. Quantification of the mean GFP pixel signal in transgenic p*_ges-1_*mtGFP animals treated with control, *plin-1, sex-1*, *sex-1(1/2)*, *plin-1(1/2)* or *plin-1;sex-1* RNAi (n=3 independent experiments, 90 animals in total, ***P < 0.001, ****P < 0.0001; Kruskal-Wallis test).

Previous interactome studies in yeast identified GTP-dependent cytosolic factors as regulators of LD–mitochondria contact sites, with the mitochondrial outer membrane protein Tom22 and the LD-associated protein Osw5 proposed as molecular tethers^14^. In *C. elegans*, TOMM-20, the orthologue of mammalian TOMM20 and a component of the mitochondrial outer membrane translocase complex, has also been identified in the LD proteome^40^. Consistent with a role in organelle contact formation, TOMM-20 protein levels increased following *sex-1* downregulation (**Figure S7A**, **Figure 4E**). To determine whether TOMM-20 contributes to the lipid storage phenotype induced by SEX-1 depletion, we simultaneously depleted SEX-1 and TOMM-20. Whereas *tomm-20* knockdown alone did not alter intestinal LD abundance, co-depletion of TOMM-20 significantly suppressed the LD accumulation induced by *sex-1* knockdown (**Figure S7B**, **Figure 4F**). These findings indicate that TOMM-20 is required, at least in part, for the effects of SEX-1 on LD accumulation, and support a role for TOMM-20 in mediating SEX-1-dependent LD–mitochondria contacts.

To further investigate the role of TOMM-20 in this inter-organelle interaction, we performed immunoprecipitation coupled to mass spectrometry (IP-MS) using p*myo-3*TOMM-20::mRFP as bait. Analysis of the TOMM-20 interactome identified 917 proteins (**Supplementary Table 5**), of which only six met both the raw *p* < 0.05 and fold-change thresholds. Among these, LET-767, a lipid-metabolizing enzyme involved in fatty acid and steroid metabolism, emerged as a top candidate. LET-767 showed an enriched pulldown trend upon *sex-1* RNAi that paralleled bait recovery, without a corresponding change in binding stoichiometry (**Figure S7C, D**). This finding is consistent with the increased LD– mitochondria contacts observed following *sex-1* depletion. Notably, LET-767, a homologue of mammalian 17β-HSD3/12, has recently been reported to localise to both LDs and the ER and to be required for the proper targeting of DHS-3 and PLIN-1 to LDs^50^.

We next investigated whether SEX-1 also regulates mitochondrial homeostasis through PLIN-1, an LD-associated perilipin homolog that is upregulated following *sex-1* downregulation (**Figure 2A**). This hypothesis is supported by studies showing that the mammalian perilipin PLIN5 promotes mitochondrial recruitment to the LD surface^15, 16^. To determine whether PLIN-1 contributes to the mitochondrial phenotype associated with SEX-1 depletion, we depleted *plin-1* in p*_ges-1_*mtGFP transgenic animals (**Figure S7E**). Similar to *sex-1* knockdown, PLIN-1 depletion significantly increased mitochondrial abundance (**Figure 4G**), indicating that LD-associated proteins contribute to mitochondrial homeostasis. However, simultaneous depletion of PLIN-1 and SEX-1 abolished the increase in mitochondrial abundance, suggesting that PLIN-1 is required for the mitochondrial phenotype induced by SEX-1 depletion (**Figure 4G**). Collectively, these findings identify TOMM-20, LET-767 and PLIN-1 as downstream effectors of SEX-1 that contribute to the regulation of LD– mitochondria contacts and organelle homeostasis.

### SEX-1 maintains AFD neuronal integrity and sensory function through a cell-autonomous mechanism

Transcriptomic analysis of *sex-1(y263)* mutants and *sex-1*::H1-wCherry transgenic animals implicated SEX-1 in neuronal development and signalling (**Figure 1C**, **D**). Analysis of published *C. elegans* ageing transcriptomic datasets^51^ further revealed that *sex-1* expression declines with age in the germline, somatic gonad, and the AFD thermosensory/CO₂-sensing amphid neurons (**Figure S8A**). To validate SEX-1expression in AFD, we generated animals co-expressing the AFD-specific reporter p*_gcy-8_*GFP and *sex-1*::H1-wCherry. Confocal microscopy confirmed SEX-1 expression in *gcy-8*-positive AFD neurons (**Figure 5A**, **B**).

**Figure 5.**
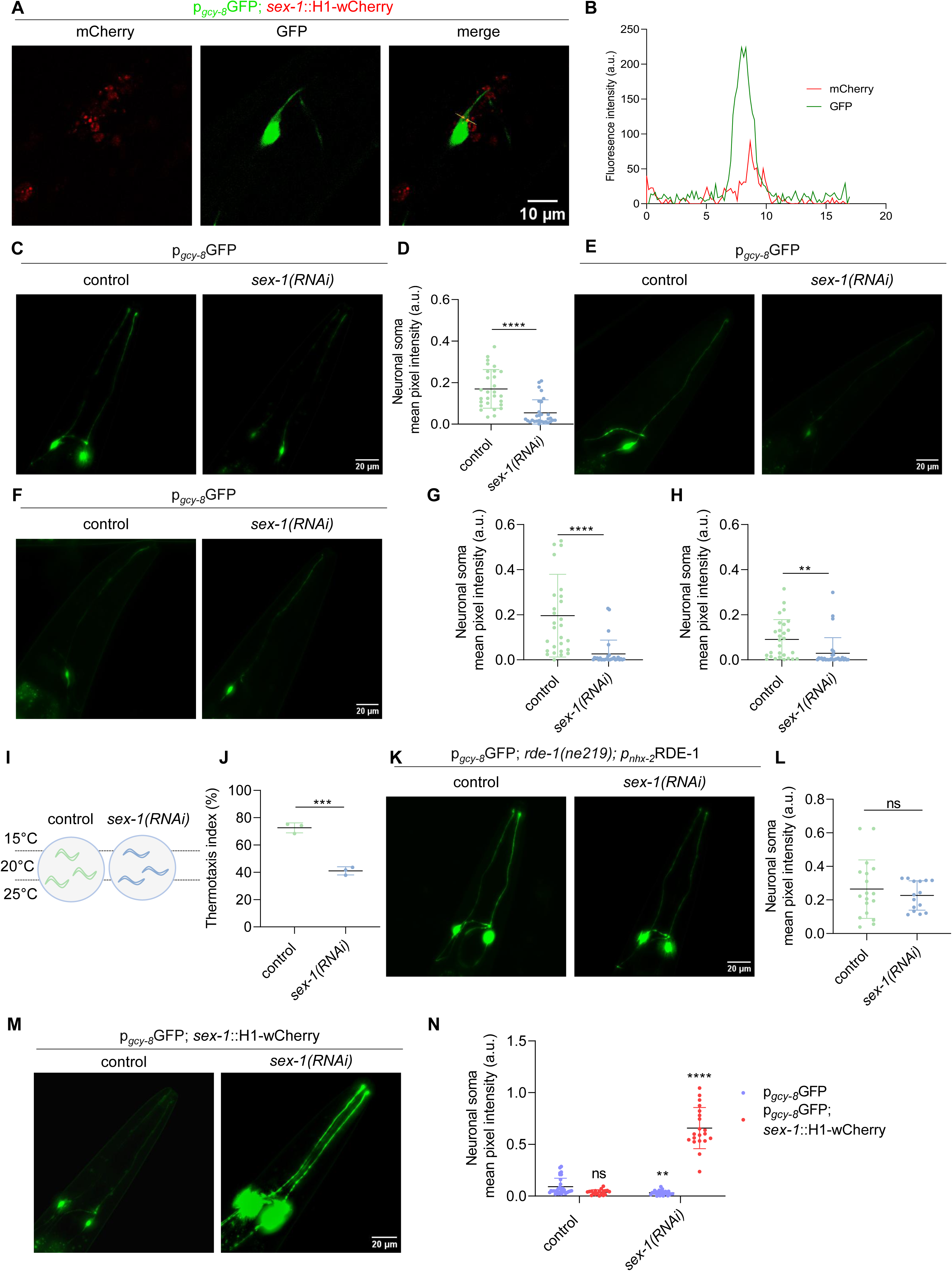
SEX-1 maintains sensory neuron integrity and function in a cell-autonomous manner. A. Representative image of transgenic animals co-expressing the *sex-1*::H1-wCherry and *gcy- 8::*GFP reporters (Images were acquired using a ×63 objective lens, n=1 experiment). B. Respective fluorescent intensity graph of the area marked with the yellow line in panel A. C. Representative images of transgenic p*_gcy-8_*GFP animals treated with control or *sex-1* RNAi showing the morphology of AFD neurons at day 1 of adulthood (Images were acquired using a ×20 objective lens, n=3 independent experiments). D. Quantification of the mean GFP signal of AFD neuronal somata as shown in (B) (n=3 independent experiments, 90 animals in total, ****P < 0.0001; two-tailed unpaired t-test). E. Representative images of transgenic p*_gcy-8_*GFP animals treated with control or *sex-1* RNAi showing the morphology of AFD neurons at day 3 of adulthood (Images were acquired using a ×4 objective lens, n=3 independent experiments). F. Representative images of transgenic p*_gcy-8_*GFP animals treated with control or *sex-1* RNAi showing the morphology of AFD neurons at day 5 of adulthood (Images were acquired using a ×4 objective lens, n=3 independent experiments). G. Quantification of the mean GFP signal in AFD neuronal somata as shown in (D) (n=3 independent experiments, 90 animals in total, ****P < 0.0001; two-tailed unpaired t-test). H. Quantification of the mean GFP signal in AFD neuronal somata as shown in (E) (n=3 independent experiments, 90 animals in total, **P < 0.01; two-tailed unpaired t-test). I. Experimental setup for monitoring thermotaxis behaviour. J. Measurement of the thermotaxis index of animals treated with control or *sex-1* RNAi (n=3 independent experiments, ***P < 0.001; two-tailed unpaired t-test). K. Representative images of transgenic p*_gcy-8_*GFP animals treated with control or *sex-1* intestine-only RNAi (Images were acquired using a ×20 objective lens, n=2 independent experiments). L. Quantification of the mean GFP signal in AFD neuronal somata as shown in (F) (n=2 independent experiments, two-tailed unpaired t-test). M. Representative images of transgenic animals co-expressing the *sex-1*::H1-wCherry and *gcy-8*::GFP reporters, treated with control or *sex-1* RNAi (Images were acquired using a ×20 objective lens, n=3 independent experiments). N. Quantification of the mean GFP signal in AFD neuronal somata as shown in (H) (n=3 independent experiments, **P < 0.01; ****P < 0.0001; Kruskal-Wallis test).

To investigate the role of SEX-1 in AFD neuronal integrity, we downregulated *sex-1* by RNAi in p*_gcy-8_*GFP transgenic animals and quantified fluorescence intensity of the AFD neuronal soma as a measure of neuronal integrity (**Figure 5C**). *sex-1* knockdown significantly reduced AFD neuronal integrity from the onset of adulthood (**Figure 5D**), and this defect persisted throughout ageing (**Figure 5E–H**). In contrast, SEX-1 depletion did not affect the morphology of other amphid neuron subtypes, including AWB, AWC and ASEL, at day 1 of adulthood (**Figure S8B-G**), indicating that SEX-1 is selectively required for maintaining AFD neuronal integrity, particularly during early adulthood.

Because the bilateral AFD neurons (AFDL and AFDR) function as the principal thermosensory neurons in *C. elegans*, capable of detecting temperature changes as small as 0.05°C, and also contribute to CO_2_ sensing and locomotory regulation^52–55^ we next examined whether SEX-1 is required for AFD sensory function. Thermotaxis assays performed on 1-day-old adults raised at 20°C demonstrated that *sex-1(RNAi)* animals exhibited impaired thermotactic behaviour compared with wild-type controls (**Figure 5I**, **J**), indicating that SEX-1 is required for normal AFD sensory function. Given that SEX-1 is expressed in AFD neurons, we investigated whether its function is cell autonomous. We generated p*_gcy-8_*GFP; *rde-1(ne219)*; p*_nhx-2_*RDE-1 transgenic animals and performed intestine-specific *sex-1* RNAi (**Figure 5K**). Intestinal depletion of SEX-1 had no detectable effect on AFD neuronal morphology (**Figure 5L**), supporting a cell-autonomous role for SEX-1 in these neurons.

To determine whether altered SEX-1 dosage influences AFD integrity, we analysed animals co-expressing p*_gcy-8_*GFP and *sex-1*::H1-wCherry. Overexpression of *sex-1* alone did not affect AFD neuronal integrity (**Figure 5M**, **N**). However, partial reduction of *sex-1* transcript levels in these double-transgenic animals significantly increased neuronal fluorescence intensity compared with control animals expressing p*_gcy-8_*GFP alone (**Figure 5M**, **N**). Together, these findings suggest that maintenance of AFD neuronal integrity requires tightly regulated *sex-1* expression levels.

### SEX-1 preserves AFD neuronal integrity by regulating sphingolipid genes *sms-5* and *mlt-8*

Having established the role of SEX-1 in lipid droplet (LD) regulation and mitochondrial homeostasis, we investigated whether its effects on AFD neuronal integrity are mediated through either pathway. To determine whether mitochondrial content is altered specifically in AFD neurons following *sex-1* knockdown, we examined transgenic animals co-expressing p*_gcy-8_*GFP and PLIN-1::mCherry, stained with MitoTracker Deep Red (**Figure S8H**). Quantification of mitochondrial colocalization with the AFD neuronal reporter revealed no significant differences between control and *sex-1(RNAi)* animals (**Figure S8I**), indicating that mitochondrial abundance in AFD neurons is not detectably affected by *sex-1* depletion.

Because our previous analyses showed that SEX-1 regulates neuronal LD abundance (**Figure 2C**), we next investigated whether altered LD homeostasis contributes to AFD degeneration. Transgenic animals co-expressing neuronal PLIN-1::GFP and *sex-1*::H1-wCherry (**Figure S9A**) demonstrated that *sex-1* overexpression significantly reduced neuronal LD abundance at day 1 of adulthood compared with age-matched PLIN-1::GFP controls (**Figure S9B**), confirming that SEX-1 negatively regulates neuronal LD accumulation. This effect was attenuated during ageing (**Figure S9C**, **D**), suggesting that SEX-1-mediated regulation of neuronal lipid homeostasis diminishes with age.

To determine whether LD accumulation occurs within AFD neurons, we generated animals co-expressing p*_gcy-8_*GFP and PLIN-1::mCherry. Although LD colocalization with the AFD marker varied between animals, *sex-1* silencing consistently increased LD abundance within AFD neurons at day 1 of adulthood (**Figure 6A**). Quantification of LD volume overlap confirmed a significant increase in AFD-associated LDs (**Figure 6B**). Thus, *sex-1* downregulation is associated with both LD accumulation and loss of AFD neuronal integrity. Consistent with the age-dependent effects of SEX-1 on neuronal LDs, differences between control and *sex-1(RNAi)* animals were no longer apparent by day 5 of adulthood (**Figure S9E**, **F**).

**Figure 6.**
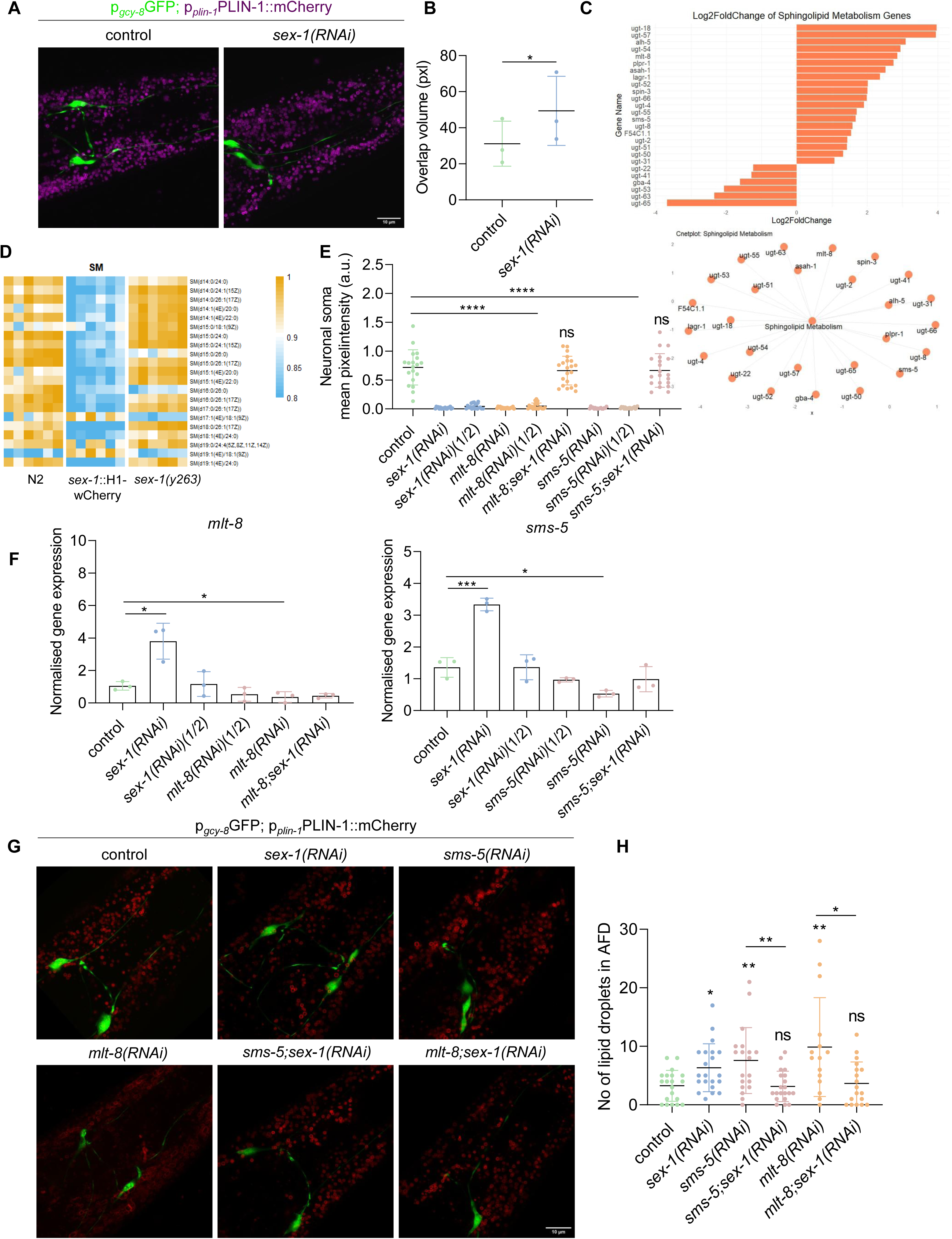
SEX-1 maintains the integrity of AFD sensory neurons by modulating the expression of sphingolipid genes *sms-5* and *mlt-8*. A. Representative single z-stack images of transgenic animals co-expressing PLIN-1::mCherry and *gcy-8*::GFP treated with control or *sex-1* RNAi (Images were acquired using a ×63 objective lens, n=3 independent experiments). B. Quantification of the overlap between lipid droplets and AFD neurons (n=3 independent experiments, *P < 0.05; two-tailed unpaired t-test). C. Bar plot and network plot from gene ontology (GO) analysis of *sex-1(y263)* mutants showing changes in expression of sphingolipid metabolism genes. The horizontal coordinate of the bar plot shows the log2fold upregulation or downregulation and the vertical coordinate shows the names of the genes. D. Heatmap comparing sphingomyelin levels in *sex-1(y263)* mutants, wild-type (N2) and *sex-1*::H1-wCherry transgenic animals. Colour intensity represents the relative changes in sphingomyelin levels, with orange indicating upregulation and blue indicating downregulation. E. Quantification of GFP signal in AFD neuronal somata of transgenic animals treated with control, *sex-1*, *sex-1(1/2), mlt-8, mlt-8(1/2)*, *mlt-8;sex-1, sms-5, sms-5(1/2)* or *sms-5;sex-1* RNAi (n=3 independent experiments,****P < 0.0001; two-tailed unpaired t-test). F. Expression analysis of *mlt-8* and *sms-5* by RT qPCR in wild-type (N2) animals treated with control, *sex-1*, *sex-1(1/2), mlt-8, mlt-8(1/2)*, *mlt-8;sex-1, sms-5, sms-5(1/2)* or *sms-5;sex-1* RNAi (n=3 independent experiments, ***P < 0.001, *P < 0.05; two-tailed unpaired t-test). G. Representative images of transgenic animals co-expressing PLIN-1::mCherry and *gcy-8*::GFP, treated with control, *sex-1*, *sms-5*, *sms-5;sex-1*, *mlt-8* or *mlt-8;sex-1* RNAi (Images were acquired using a ×63 objective lens, n=3 independent experiments). H. Number of lipid droplets in AFD neurons of transgenic animals shown in (G), treated with control, *sex-1*, *sms-5*, *sms-5;sex-1*, *mlt-8* or *mlt-8;sex-1* RNAi (n=3 independent experiments,*P < 0.05, **P < 0.01; Mann-Whitney test).

Aberrant LD accumulation has been linked to dysregulated sphingolipid metabolism^56^. Consistent with this relationship, transcriptomic and lipidomic analyses of *sex-1(y263)* mutants revealed increased expression of genes involved in sphingolipid metabolism together with elevated sphingomyelin levels (**Figure 6C**, **D**). To identify modifiers of AFD degeneration, we performed an RNAi screen in p*_gcy-8_*GFP animals targeting genes that were upregulated more than twofold in *sex-1(y263)* mutants relative to wild-type animals. This screen identified *sms-5* and *mlt-8*, encoding a sphingomyelin synthase and a molting protein implicated in glycosphingolipid catabolism, respectively, as modifiers of AFD degeneration. SMS-5 localizes to the Golgi apparatus and plasma membrane, where it catalyses sphingomyelin production from ceramide and phosphatidylcholine while generating diacylglycerol^57^. MLT-8 localizes to lysosomes and is subsequently secreted to the cuticle^58^.

Unexpectedly, knockdown of either *sms-5* or *mlt-8* alone impaired AFD neuronal integrity, whereas simultaneous knockdown of either gene together with *sex-1* rescued the degeneration phenotype (**Figure 6E**, **Figure S9G**), indicating a genetic interaction between *sex-1* and these sphingolipid regulators. Consistent with this model, *sex-1* knockdown significantly increased the expression of both *sms-5* and *mlt-8*, whereas this induction was suppressed in *sms-5(RNAi)*;*sex-1(RNAi)* and *mlt-8(RNAi)*;*sex-1(RNAi)* animals (**Figure 6F**).

Finally, super-resolution confocal microscopy of p*_gcy-8_*GFP;PLIN-1::mCherry animals revealed that depletion of either *sms-5* or *mlt-8* increased LD abundance within AFD neurons (**Figure 6G**, **H**). In contrast, simultaneous knockdown of *sex-1* significantly reduced LD accumulation in both genetic backgrounds (**Figure 6G, H**). Together, these findings indicate that SEX-1 preserves AFD neuronal integrity by regulating the expression of *sms-5* and *mlt-8*, thereby maintaining sphingolipid-dependent lipid droplet homeostasis.

## Discussion

Our findings identify SEX-1 as a functional orthologue of human PPARγ and establish a previously unrecognised role for PPARγ-like signalling in coordinating lipid homeostasis across lipid droplets (LDs) and mitochondria while preserving neuronal integrity. Pharmacological activation with rosiglitazone further supports the utility of *C. elegans* as a genetically tractable model for investigating PPARγ regulation and function. We demonstrate that SEX-1 regulates LD abundance and size, mitochondrial mass and function, and LD–mitochondria interactions across multiple tissues, while maintaining the structural and functional integrity of AFD thermosensory neurons. Transcriptomic analyses further implicate SEX-1 in the regulation of sphingolipid-associated pathways, including *mlt-8* and *sms-5*, thereby linking sphingolipid homeostasis to neuronal LD dynamics. Collectively, these findings define a PPARγ-like signalling network that integrates lipid metabolism, organelle homeostasis, and neuronal function, providing mechanistic insight into how metabolic homeostasis supports neuroprotection (**Figure 7**).

**Figure 7.**
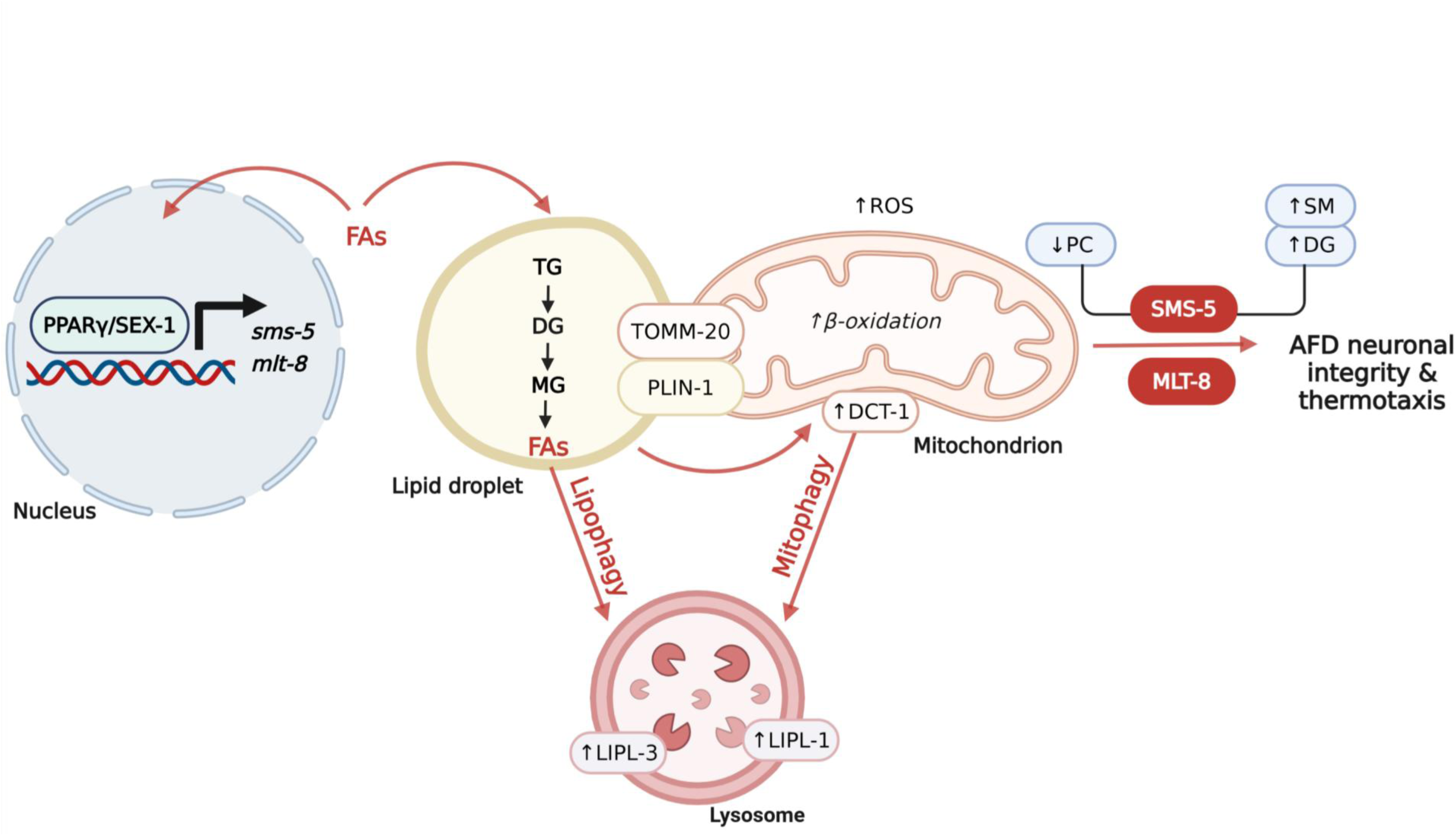
Schematic representation of SEX-1/ PPARγ-mediated modulation of systemic metabolism and neuronal integrity. SEX-1/ PPARγ modulates the expression of several lipid metabolism-related genes, such as *sms-5* and *mlt-8*, thereby influencing the abundance of various lipid species, including sphingomyelin (SM), phosphatidylcholine (PC), and diacylglycerol (DG). As endogenous ligands for PPARs, fatty acids (FAs), may either activate SEX-1/ PPARγ or be sequestered into lipid droplets (LDs) for storage as triacylglycerols (TG). SEX-1/ PPARγ regulates TG storage by modulating LD number and size, partly through sterol regulatory element-binding protein (SBP-1)-mediated lipogenesis and lipophagy. In addition, SEX-1 influences inter-organelle contacts, particularly between LDs and mitochondria, potentially facilitating FAs transfer for β-oxidation and contributing to elevated reactive oxygen species (ROS) production. SEX-1/ PPARγ also modulates mitochondrial mass by coordinating mitochondrial quality control pathways associated with DCT-1-dependent mitophagy, while influencing the abundance of potential inter-organelle interactors, such as PLIN-1 and TOMM-20. Finally, SEX-1/ PPARγ plays a crucial role in neuronal integrity, particularly in AFD thermosensory neurons, by regulating neuronal LD accumulation and sphingolipid-associated pathways linked to *sms-5* and *mlt-8*, which in turn influence DG and SM abundance (created with BioRender.com).

Consistent with the established role of PPARγ in mammalian lipid metabolism ^8, 59^, SEX-1 emerged as a key regulator of LD homeostasis. Overexpression of *sex-1* reduced LD accumulation, whereas *sex-1* depletion increased both LD number and size across multiple tissues and was accompanied by elevated triglyceride and sphingomyelin levels. Mechanistically, LD accumulation in SEX-1-deficient animals resulted from enhanced SBP-1-dependent lipogenesis together with activation of lipophagy, rather than increased lipolysis. The absence of detectable lipolysis despite lipid accumulation may reflect the relatively modest increase in LD size, which remains below the threshold proposed to efficiently stimulate lipolysis^60^. These findings extend the emerging role of PPARγ beyond its established functions in lipid synthesis and degradation^36, 59^, by implicating it in the regulation of lipophagy, although the underlying molecular mechanisms remain poorly understood^61, 62^. Whether SEX-1 regulates lipophagy through direct modulation of lysosomal lipases, including *lipl-1* and *lipl-3*, or indirectly as a compensatory response to elevated lipogenesis remains to be determined. Nevertheless, our data support the concept that PPARγ-like signalling acts as a central metabolic hub integrating lipogenic and autophagic pathways to maintain lipid homeostasis.

The close functional relationship between LDs and mitochondria is increasingly recognised as a major determinant of cellular lipid metabolism^12^. In mammalian cells, PPARγ activation promotes mitochondrial biogenesis and fatty acid β-oxidation, primarily through the PGC1α pathway^63–65^. In contrast, *sex-1* downregulation increased mitochondrial mass in *C. elegans*, suggesting divergence in the mechanisms governing mitochondrial homeostasis. As *C. elegans* lacks a clear PGC1α orthologue, alternative regulators such as SKN-1/Nrf2, which coordinates mitochondrial stress responses and function, may mediate these effects^48, 66^. Loss of SEX-1 resulted in mitochondrial hyperactivity characterised by elevated oxygen consumption and reactive oxygen species (ROS) production, accompanied by increased mitophagy despite minimal alterations in mitochondrial morphology. Reduced ATP levels further indicate that increased respiratory activity is inefficiently coupled to ATP synthesis^12, 47^, consistent with mitochondrial dysfunction rather than enhanced bioenergetic capacity. In parallel, *sir-2.2*, a gene encoding a mitochondrial sirtuin involved in fatty acid oxidation and regulation of mitochondrial gene expression^49, 67^, was downregulated following *sex-1* knockdown, and *sir-2.2* depletion phenocopied key mitochondrial defects observed in SEX-1-deficient animals. Together, these findings identify SEX-1 as an important regulator of mitochondrial quality control, coordinating mitochondrial turnover and function in response to altered lipid metabolism.

Our study also reveals an unexpected role for SEX-1 in regulating LD–mitochondria interactions. Although bidirectional communication between these organelles is well established^15, 16, 68^, the involvement of PPARγ signalling in this process has not previously been demonstrated. Loss of *sex-1* reduced the distance between LDs and mitochondria, indicating tighter physical association that coincided with increased triglyceride accumulation and LD expansion. This enhanced proximity was accompanied by mitochondrial hyperactivity, consistent with previous observations that LD-associated mitochondria display elevated oxidative metabolism^15, 69^. Our data further identify TOMM-20, LET-767 and PLIN-1 as candidate mediators of this organelle interface. Notably, depletion of PLIN-1, which encodes a key regulator of LD–mitochondria interactions^10^, caused an even greater increase in mitochondrial mass than *sex-1* knockdown alone. This observation is consistent with studies showing increased mitochondrial abundance in the livers of Plin5-deficient mice^70^, and supports a role for perilipins in regulating mitochondrial homeostasis. Although the molecular architecture underlying LD– mitochondria tethering remains incompletely understood, our findings position SEX-1/PPARγ signalling as an important upstream regulator of organelle crosstalk and metabolic coordination.

Beyond its metabolic functions, our data establish SEX-1 as a critical regulator of neuronal homeostasis. Loss of SEX-1 induced mitochondrial hyperactivity, increased ROS production, and neuronal LD accumulation, all hallmarks commonly associated with neurodegeneration^43, 71, 72^. Importantly, SEX-1 depletion compromised both the structural integrity and thermotactic function of AFD neurons from early adulthood, demonstrating a direct requirement for SEX-1 in maintaining neuronal resilience. Transcriptomic analyses identified increased expression of *mlt-8* and *sms-5*, genes associated with sphingolipid metabolism, as potential mediators of these defects. Partial suppression of either gene attenuated both neuronal degeneration and LD accumulation in AFD neurons, indicating that dysregulated sphingolipid metabolism contributes to the neuronal phenotypes associated with loss of SEX-1. Under physiological conditions, neurons contain relatively few LDs because excess lipids are continuously transferred to glia and fatty acids are rapidly metabolised. Consequently, neuronal LD accumulation following SEX-1 depletion is likely to reflect a protective response that buffers lipotoxicity and oxidative stress rather than a primary pathological event. These findings therefore establish a mechanistic link between PPARγ-like signalling, sphingolipid metabolism, organelle dynamics, and neuronal maintenance.

In summary, our study identifies an unexpected role for the *C. elegans* PPARγ orthologue SEX-1 in coordinating metabolic and neuronal programmes, thereby expanding the functional repertoire of PPARγ-like signalling beyond its established roles in lipid and glucose metabolism. By integrating sphingolipid metabolism, LD dynamics, mitochondrial quality control, and neuronal integrity, our findings reveal a broader framework through which PPARγ-like pathways coordinate cellular adaptation to metabolic stress. Several important questions remain. The molecular mechanisms through which SEX-1 regulates LD–mitochondria interactions require further investigation, as does the extent to which analogous pathways operate in mammalian neurons. Moreover, given the tissue-specific and context-dependent functions of PPARγ isoforms, caution is warranted when extrapolating findings from *C. elegans* directly to human physiology. Nevertheless, the identification of a neuronal– metabolic regulatory axis governed by a PPARγ orthologue provides a conceptual framework for understanding how metabolic signalling integrates organelle homeostasis with neuronal maintenance, with important implications for ageing, neurodegeneration, and metabolic disease.

## Materials and Methods

### C. elegans strains

Standard procedures were followed for maintaining *C. elegans* strains. The rearing temperature was set at 20°C unless otherwise noted. All the strains used in this study are listed in **Supplementary Table 6**.

### RNAi and molecular cloning

For knockdown of genes by RNAi, we used HT115 (DE3) *E. coli* bacteria transformed with the L4440 plasmid vector expressing double-stranded RNA against the gene of interest. The HT115 strains were either obtained from the Ahringer library or RNAi constructs were generated by PCR amplification of gene-specific regions using *C. elegans* genomic DNA as a template and primer sets as annotated in **Supplementary Table 7**. Thereafter, the generated constructs were inserted into the TOPO-pCRII vector (Invitrogen) by TA cloning and sub-cloned into the final pL4440 vector. Positive clones were transformed into HT115 bacteria to allow knockdown of the genes of interest through feeding^73^. For *sex-1* RNAi, we amplified a 1,713 bp region using *sex-1* RNAi Forward and *sex-1* RNAi Reverse primers. For *mlt-8* and *sms-5* RNAi, the Ahringer *C. elegans* RNAi feeding library was used. To conduct RNAi experiments, a single bacterial colony was used to inoculate LB medium containing 100 µg/ml ampicillin and 10 µg/ml tetracycline (**Supplementary Table 8**) overnight at 37°C in a shaker with 150 rpm for 16 hours. Thereafter, 250 µl of overnight culture were inoculated in LB medium containing 100 µg/ml ampicillin for 3-4 hours and RNAi plates were seeded with 250 µl of bacterial culture and 2 mM isopropylthiogalactoside (IPTG). RNAi plates were used the next day and were always freshly prepared.

### Docking of Rosiglitazone to SEX-1

The predicted three-dimensional structures of the *C. elegans* transcription factor SEX-1 and human PPARγ were obtained using AlphaFold, in PDB formats. The structures were prepared for docking using the (mk_prepare_receptor.py) script from AutoDock Tools, with water molecules removed (-- delete_residues HOH) and problematic residues automatically skipped using the (--allow_bad_res) flag. The ligand rosiglitazone was converted to pdbqt format using standard ligand preparation protocols. The docking search box was generated by enclosing the ligand with a 4 Å padding (-- box_enveloping rosiglitazone.pdbqt --padding 4). Molecular docking was carried out using AutoDock Vina v1.2.3^28, 29^, generating multiple ligand poses based on predicted binding affinity. The docking scores ranged from –∼6 to –∼2 kcal/mol, with the top pose showing the strongest predicted interaction. The results were analyzed using the ViewDockX plugin in UCSF ChimeraX v1.9, which enabled clustering based on RMSD, visualization of binding poses, and identification of potential interaction residues within the binding pocket.

### Compound treatment

Rosiglitazone (**Supplementary Table 8**) was added to the NGM at three different final concentrations: 0.1, 1, 10 µM, or 1% DMSO (**Supplementary Table 8**), as previously described^74^. *E. coli* OP50 bacteria were heat-inactivated at 65°C for 45 minutes to avoid interference by the xenobiotic metabolizing activity and diluted in S basal buffer. Plates were allowed to dry for one day and always prepared fresh. Animals were synchronised by bleaching and grown on rosiglitazone plates from eggs to day 1 of adulthood.

### mRNA quantification and qPCR

To quantify gene expression, total RNA was extracted using the TRIzol reagent (Invitrogen). All the primers used are summarized in **Supplementary Table 7**. For cDNA synthesis, mRNA was reverse transcribed using an iScriptTM cDNA Synthesis Kit (BioRad). Quantitative PCR was performed in triplicate with the Eva Green qPCR Kit (Biotium) in a Bio-Rad CFX96 Real-Time PCR system (Bio-Rad), according to manufacturer’s instructions. The following housekeeping genes were used: *pmp-3*, *actin-1, rps-23,* or *ama-1*.

### RNA sequencing and data analysis

To examine possible targets of SEX-1, we performed a wide transcriptomic analysis on N2, *sex- 1(y263)* mutants and *sex-1*::H1-wCherry transgenic animals at day 1 of adulthood. For each strain, five biological replicates were obtained. Total RNA was extracted using the TRIzol reagent and RNA sequencing was performed by Alithea Genomics, as well as the bioinformatic analysis and on average 21 million reads were obtained for each sample.

### Whole genome sequencing

To genetically characterize the *sex-1*::H1-wCherry transgenic animals, genomic DNA was extracted from a mixed-stage population. High molecular weight DNA was prepared and submitted for long-read sequencing using the Oxford Nanopore platform. The bioinformatic analysis of the plasmid sequence and integration events was performed by the Bioinformatics Unit at the Institute of Molecular Biology and Biotechnology (IMBB-FORTH). Transgene integration sites were identified by searching for contigs that aligned to both chromosomal DNA and the transgene plasmid.

### Live Imaging

Confocal images of adult worms were acquired with LEICA TCS SP8 laser scanning confocal microscope or ZEISS LSM 900 with Airyscan, using x40 or x63 objective lens. For epifluorescence microscopy, images were acquired with EVOS FL Auto 2 imaging system (AMAFD2000; Thermo Fisher Scientific). In all cases, animals were anesthetized using 20 mM levamisole (**Supplementary Table 8**) before mounting and image analysis was performed in Fiji version 2.0.0.

### Confocal microscopy for lipid droplet quantification

To quantify the size and number of lipid droplets by super-resolution confocal microscopy, we used three different reporter strains expressing lipid droplet proteins: DHS-3 fused to GFP, which is expressed only in the intestine, PLIN-1 fused to mCherry, which is expressed ubiquitously, and PLIN1 fused to GFP, expressed pan-neuronally. The first two transgenic animals (LIU1, LIU2) were imaged at day 1 of adulthood, while *xdIs109* animals were imaged at day 1 and 5 of adulthood. For each experiment, 30 worms were imaged in total, per condition, anesthetized with 20 mM levamisole and covered with a glass coverslip for imaging. The images in each experiment were taken using a x63 objective lens, with the same zoom, pinhole and exposure time/laser power. The number and size of all lipid droplets in focus were analyzed in Fiji version 2.0.0 manually in the same area of the intestine or the head.

### Imaging of mitochondrial network

We used two different reporter strains: SJ4103 and SJ4143 that express GFP in the matrix of mitochondria in the body wall muscle and intestine, respectively. To measure mitochondrial mass, mean fluorescent intensity was measured in whole animals using epifluorescence microscopy. To measure mitochondrial morphology, Z stack images were acquired using confocal microscopy with a x63 objective lens and the mean branch length and mitochondrial network branches were quantified in Fiji version 2.0.0 using the plugin MiNA.

### Confocal microscopy for lipid droplet-mitochondria interaction

Acquired images were analysed using the ImageJ plugin Diana (surface contact analysis). Lipid droplets (first image) were segmented using the classic segmentation method with no filter and the mitochondria were segmented using the same method but with a Gaussian filter and an XY radius of 0.5. In both cases, the minimum and maximum object size was set at 3 and 20,000 pixels, respectively. The same threshold was used for all the images of each experiment. DiAna colocalization analysis was performed to calculate the surface in contact length between lipid droplets and mitochondria. The maximum distance to establish contact between objects was set at 50 nm.

### Immunoprecipitation–mass spectrometry

∼5000 synchronized PS6187 transgenic animals, treated with control or *sex-1* RNAi, were collected at day 1 of adulthood in two independent experiments. *unc-119(ed3)* mutants were also included as a negative control for nonspecific protein binding. Worms were collected in M9 buffer, washed five times, frozen in liquid nitrogen and stored at -80°C. For protein extraction, worm pellets were lysed in 0.2 ml lysis buffer (50mM Tris pH 8.0, 150mM NaCl, 1% NP-40, 1mM MgCl_2_, 1 mM EDTA, 1 mM EGTA, 1 mM PMSF, Protease Inhibitor Cocktail (Sigma-Aldrich)), by five freeze-thaw cycles, followed by three cycles of sonication. For immunoprecipitation, 15 µl magnetic beads were incubated with 1.5 µg goat polyclonal anti-RFP antibody (**Supplementary Table 8**) or without antibody as a negative control. Immunoprecipitated samples were analyzed using a Vanquish Neo UHPLC system (Thermo Fisher Scientific) operated in trap-and-elute mode and coupled online to an Orbitrap Excedion Pro mass spectrometer (Thermo Fisher Scientific). Mass spectrometry was performed in the Proteomics Facility at Institute of Molecular Biology and Biotechnology, Foundation for Research and Technology - Hellas (PROFI). TOMM-20::RFP interactors, were identified by comparison of positive samples with both *unc-119(ed3)* and no-antibody controls.

### Sample preparation for lipidomics

To analyse lipid composition using mass spectrometry, wild-type, *sex-1(y263)* mutants and *sex-1*::H1-wCherry transgenic animals were collected at day 1 of adulthood in six technical replicates. Synchronized animals were collected in M9 buffer in 15 ml tubes and washed at least four times, by allowing the worms to settle by gravity between washes to remove bacteria and eggs. Finally, worms were transferred to 1.5 ml tubes (Eppendorf), frozen with liquid nitrogen and stored at -80°C. A 5 mm steel bead and 1:1 (v/v) methanol:chloroform was added to each sample to a volume of 1.5 ml. Samples were homogenized using a TissueLyser II (Qiagen) for 5 min at 30 Hz and centrifuged for 10 min at 20,000 g. The protein concentration of the lysate was determined using a Pierce BCA protein assay kit (Thermo Scientific).

### Lipidomics analysis

The lipidomics analysis was performed as previously described^75, 76^. Mixed internal standards were added to correct for variation in lipid extraction and analytical procedures. Following extraction, the supernatant was transferred to 2 mL tubes and evaporated to dryness at 45°C. The dried lipid residues were reconstituted in 50 μL of chloroform/methanol/water (60:30:4.5, v/v/v) and subsequently diluted with 150 μL of isopropanol/acetonitrile/water (2:1:1, v/v/v). Lipid separation was performed using an Acquity UPLC system (Waters, Manchester, UK) equipped with an Acquity UPLC CSH column (1.7 μm, 100 × 2.1 mm). The mobile phase consisted of 10 mM ammonium formate in water (eluent A) and 10 mM ammonium formate in methanol (eluent B). The gradient elution program was: 0–5 min, 50–30% A; 5–15 min, 30–10% A; 15–25 min, 10–0% A, followed by isocratic elution at 0% A for 15 min. A 5-min conditioning cycle with initial mobile phase proportions was performed between analyses. The column temperature was maintained at 80°C with a flow rate of 0.5 mL/min. Sample injection volumes were 4 μL and 8 μL for positive and negative ionization modes, respectively. Mass spectrometry was performed using a Synapt G2-Si high-resolution quadrupole time-of-flight (QTOF) mass spectrometer (Waters, Manchester, UK). Nitrogen was used as the desolvation gas and argon as the collision gas. Data acquisition was performed in continuum and enhanced resolution modes over an m/z range of 50–1750 Da at an acquisition rate of 1 spectrum per 0.2 s. Source parameters were optimized as follows: source temperature, 150°C; desolvation temperature, 400°C; cone voltage, 30 V; capillary voltage, 2000 V. MS experiments were conducted in MS mode with LockSpray internal reference to ensure high mass accuracy throughout the analysis.

### Oil Red O staining

To measure fat accumulation, synchronized adult animals were stained with Oil Red O (**Supplementary Table 8**) that stains only neutral lipids^39^. Briefly, 100 worms were washed three times with M9 and allowed to settle at the bottom by gravity. The supernatant was removed, and worms were fixed with 1% paraformaldehyde (PFA), dissolved in M9 at a 1:1 ratio. The samples were left to rotate for 5 min at room temperature (RT). They were then freeze-thawed three times and washed again with M9 three times to remove PFA. The worms were then resuspended in 60% isopropanol and incubated for 15 min at room temperature. After the worms were allowed to settle at the bottom, the isopropanol was removed, 1 mL of 60% Oil-Red-O stain was added and the animals were incubated at RT for 4 hours. The stain was removed by washing three times with M9. Animals were mounted on slides and imaged using an EVOS FL Auto 2 imaging system (AMAFD2000; Thermo Fisher Scientific).

### MitoTracker CM-H2X ROS/ MitoTracker Deep Red/ TMRE staining

To assess mitochondrial ROS, mitochondrial content and mitochondrial function, animals were stained with MitoTracker Red CM-H2XRos, MitoTracker Deep Red or TMRE (tetramethylrhodamine, ethyl ester, perchlorate; a dye that accumulates in intact mitochondria) (**Supplementary Table 8**). Briefly, synchronized 1-day-old animals were transferred at the L4 stage on plates containing heat-inactivated RNAi bacteria (UV, 15 min) and 100 µl of MitoTracker or TMRE at a final concentration of 0.1 µM and 0.05 µM, respectively. Animals were incubated at 20°C overnight before immobilization with 20mM levamisole and mounting for microscopic examination using an EVOS FL Auto 2 imaging system (AMAFD2000; Thermo Fisher Scientific).

### ATP measurements

To determine ATP content, 100 synchronized 1-day-old worms were collected in M9 buffer and washed three times in three biological replicates. Supernatant was removed and pellet was frozen at - 80°C. Frozen worms were sonicated using three repetitions of 60 watt for 10 sec, per sample. 50 µl of luciferase was added to each sample just prior to measurement in a TD-20/20 luminometer (Turner Designs) and results were compared to a standard curve. ATP levels were determined by using the ATP bioluminescence assay kit CLS II (Roche Applied Science, 11699695001) and normalized to total protein content.

### Oxygen consumption rate measurements

Oxygen consumption was assessed using a Bioscience Seahorse XF96 Flux Analyser (or XFe96) as previously described^77^. Synchronized 1-day-old worms were collected in M9 buffer and washed several times to remove bacteria and larvae. Each experiment was repeated three times.

### Bioinformatic tools

Multiple sequence alignment was performed using CLUSTALW with default gap opening penalty (10.0) and gap extension penalty (0.2). The alignment was visualized and analysed using Jalview v2.0^78^. Jalview was also used to generate the figure shown in this study, to highlight sequence conservation. Structural alignments were performed using FATCAT v2.0 with default parameters to compare protein 3D structure^26^. Phylogenetic analysis was performed using MEGA 11 software^79^. The evolutionary history was inferred using the Maximum Likelihood method based on the JTT matrix-based model^80^. The initial trees for heuristic search were generated using Neighbor-Join and BioNJ algorithms, with the best topology selected based on log likelihood. A discrete Gamma distribution (5 categories, +G, parameter = 7.5020) modelled rate variation among sites, and a proportion of sites were assumed to be invariable (+I, 4.43%). The final analysis included 8 amino acid sequences with 666 aligned positions. The tree with the highest log likelihood (–10443.85) is shown, with branch support values indicating the percentage of trees in which associated taxa clustered together.

### Statistical analysis

We used the Prism software package (GraphPad Software 8) for statistical analysis. The statistical tests applied for each experiment are specified in the Figure legends. Error bars in all figures represent standard deviations (SD).

## Acknowledgements

We thank Eleftherios Morres for his valuable support with the bioinformatics analyses, as well as the IMBB Bioinformatics Unit, IMBB-FORTH, Heraklion, Crete, Greece. We thank the Proteomics Facility at IMBB-FORTH (PROFI) for assistance with sample preparation, protocol optimisation and mass spectrometry analysis. We also thank the *Caenorhabditis* Genetics Center (CGC), which is funded by the NIH Office of Research Infrastructure Programs (P40 OD010440), for providing some of the nematode strains used here. Work in the authors’ laboratory is funded by the Hellenic Foundation for Research and Innovation (H.F.R.I.) under the “1st Call for H.F.R.I. Research Projects to support Faculty members and Researchers and the procurement of high-cost research equipment” (Project Number: HFRI—FM17C3-0869, NeuroMitophagy), the H.F.R.I. Project Number:15546, Acronym: NeuroFlame, the General Secretariat for Research and Innovation of the Greek Ministry of Development, the European Union – NextGenerationEU (project code: TAEDR-0535850, Acronym: BrainPrecision) within the framework of the Action “Flagship Actions in Interdisciplinary Scientific Areas with Special Interest in Connection to the Productive Fabric,” under the National Recovery and Resilience Plan “Greece 2.0” and the European Commission Research Executive Agency Excellence Hub “CHAngeing” (GA-101087071). D. T. is supported by the Hellenic Foundation for Research and Innovation (H.F.R.I.) [Scholarship Code: 11403]. T.Z. is supported by CSC fellowship and E.N. by ParkinsonFonds NL.

## Author contributions

D.T.: Conceptualization; resources; formal analysis; validation; investigation; visualization; methodology; writing – original draft. M. M.: Conceptualization; formal analysis; supervision; investigation; methodology; writing – original draft; writing – review and editing. T. Z.: Lipidomics experiments and analysis; ORO experiments; oxygen consumption experiments. E. N.: Review and editing. N. T.: Conceptualization; resources; formal analysis; supervision; funding acquisition; writing – original draft; writing – review and editing.

## Data availability

The RNA sequencing data that support the findings of this study have been deposited in the Gene Expression Omnibus (GEO) under the accession number GSE300054

## Supplementary Figure legends

**Supplementary Figure 1.**
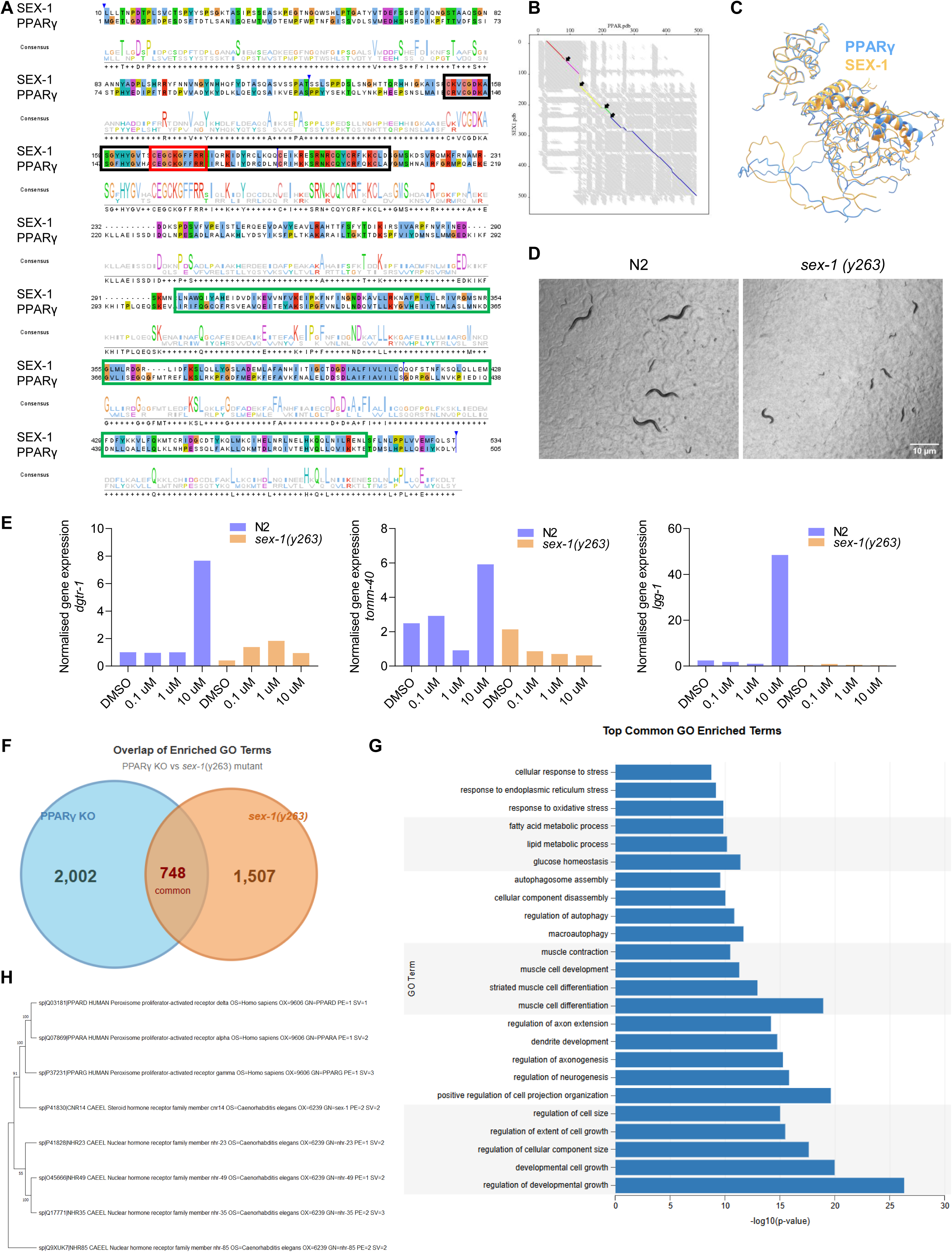
Characterisation of SEX-1, a putative *C. elegans* orthologue of PPARγ. A. CLUSTAL W multiple sequence alignment between human PPARγ (P37231.3) and nematode SEX-1 (NP_001024662.1) showing conserved regions. Amino acids are coloured according to their physicochemical properties and no colour means low conservation. The coloured areas show the conserved domains (DNA-binding domain – black, ligand-binding domain – green, P-box motif – red). B. The chaining result from flexible FATCAT alignment showing the aligned fragment pairs in the optimal alignment shown as coloured lines and all the aligned fragment pairs (AFPs) between the two structures shown as short grey lines in the background. C. The 3D visualization of superposition between PPARγ and SEX-1 proteins acquired by flexible FATCAT alignment (blue: PPARγ, yellow: SEX-1). D. Representative bright field images of wild-type (N2) and *sex-1(y263)* mutants showing the dumpy and egg laying defective phenotype. E. Expression analysis of *dgtr-1*, *tomm-40* and *lgg-1* by RT qPCR in wild-type (N2) animals and *sex-1(y263)* mutants treated with 1% DMSO or 0.1,1,10 µM Rosiglitazone (n=1 experiment). F. Venn diagram showing the overlap of enriched gene ontology (GO) terms in *sex-1(y263)* mutants and PPARγ-deficient multipotent mesenchymal stromal cells. Numbers indicate terms enriched only in each transcription factor (non-overlapping regions) or co-regulated by both (overlapping region). G. Bar plot of the top common gene ontology (GO) enriched terms in *sex-1(y263)* mutants and PPARγ-deficient multipotent mesenchymal stromal cells. The horizontal coordinate of the bar plot represents statistical significance (-log₁₀ p-value) and the vertical coordinate shows the names of the enriched terms. H. Phylogenetic tree developed using MEGA 11.0 software. Eight protein sequences were aligned with CLUSTAL W. The alignment was used to construct a tree using the Maximum Likelihood method and a Jones-Taylor-Thornton (JTT) matrix-based model, p-distance and 100 Bootstrap repetitions. The numbers at the branch points show the bootstrap values.

**Supplementary Figure 2.**
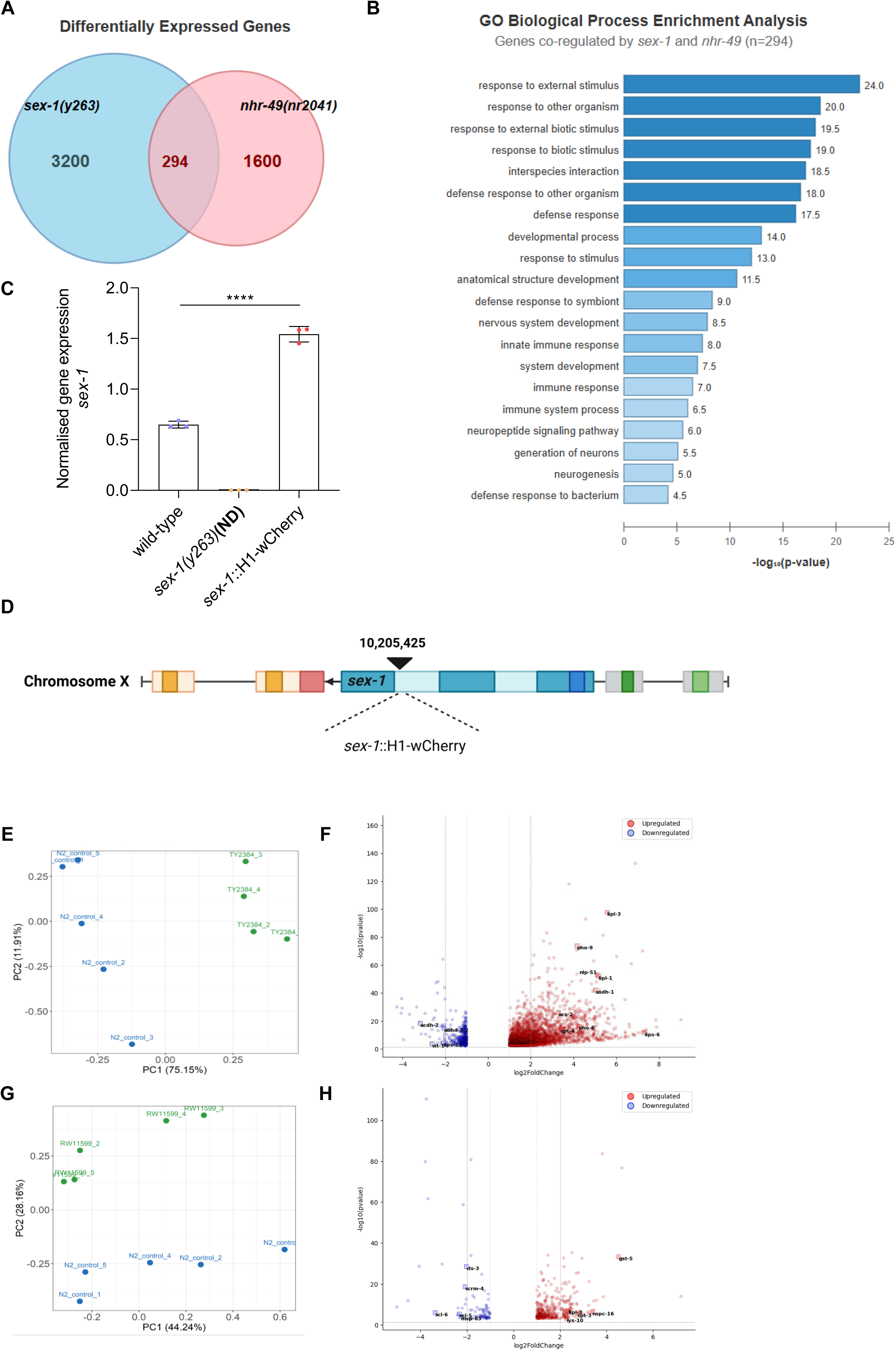
Transcriptional profiling of SEX-1. A. Venn diagram showing the overlap of differentially expressed genes (DEGs) between *sex- 1(y263)* and *nhr-49(nr2041)* mutants. Numbers indicate terms enriched only in each transcription factor (non-overlapping regions) or co-regulated by both (overlapping region). B. Bar plot of gene ontology (GO) biological process enrichment analysis of 294 genes co-regulated by *sex-1* and *nhr-49*. Bar length represents statistical significance (-log₁₀ p-value). C. Expression analysis of *sex-1* by RT qPCR in wild-type (N2) animals, *sex-1(y263)* mutants and *sex-1*::H1-wCherry transgenic animals (n=3 experiments, ****P < 0.0001; two-tailed unpaired t-test, ND = not detected after 40 cycles) D. Schematic representation of the transgene integration site identified by whole-genome sequencing in *sex-1*::H1-wCherry transgenic animals. Black triangle indicates integration point with precise genomic coordinate shown (Created with BioRender.com). E. Principal component analysis (PCA) plot of transcriptome analysis showing that N2 and TY2384 [*sex-1(y263)*] animals were grouped in different clusters. F. Volcano plot of gene expression changes in *sex-1(y263)* mutants compared to wild-type (N2) animals. Differentially expressed genes (DEGs) that are upregulated are highlighted as red and the ones that are downregulated are marked as blue (p value <0.05). G. Principal component analysis (PCA) plot of transcriptome data showing that N2 and RW11599 (*sex-1*::H1-wCherry) animals were grouped in different clusters. H. Volcano plot of gene expression changes in *sex-1*::H1-wCherry animals compared to wild-type (N2) animals. Differentially expressed genes (DEGs) that are upregulated are highlighted as red and the ones that are downregulated are marked as blue (p value < 0.05)

**Supplementary Figure 3.**
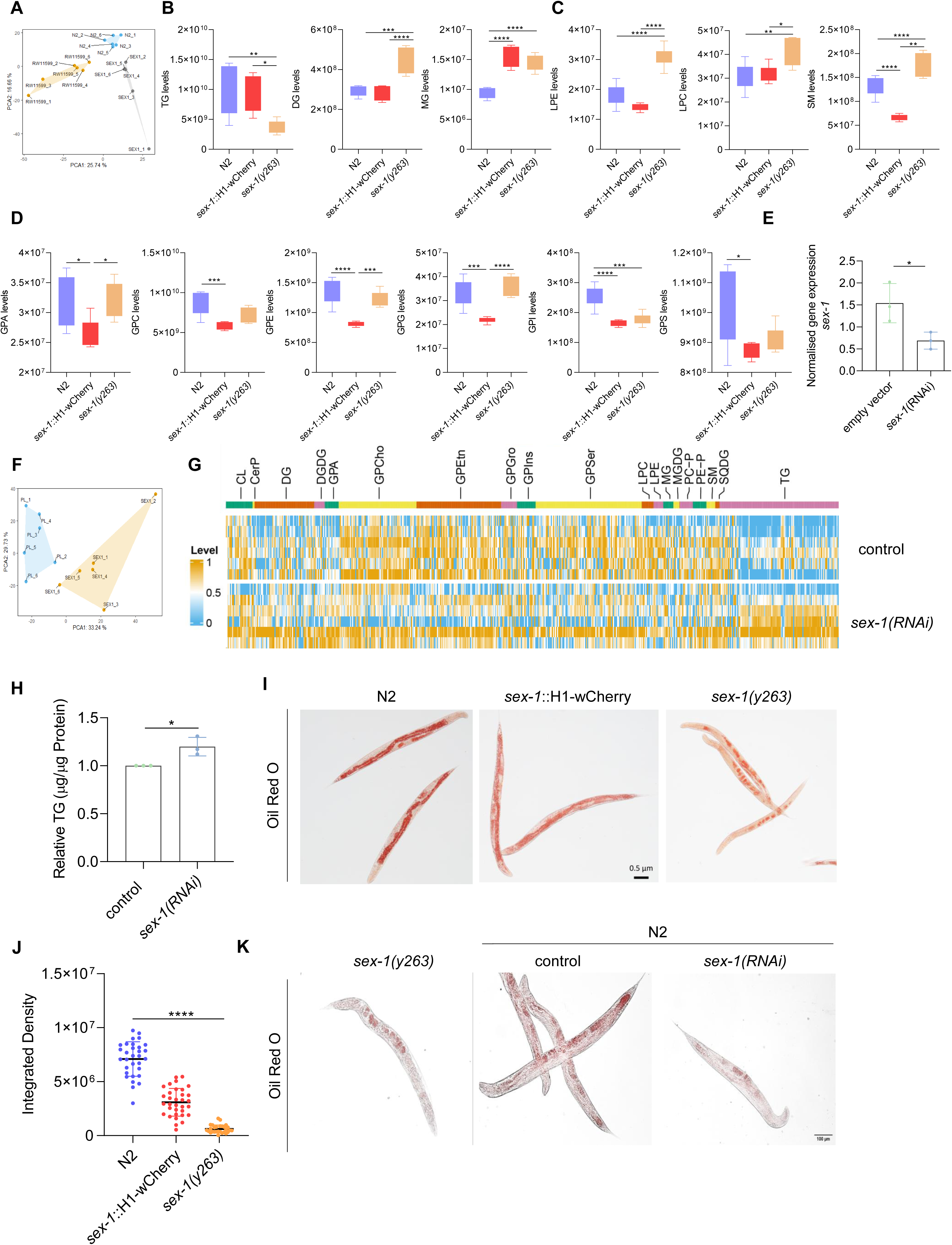
Lipidomic profiles of *sex-1(y263)* mutants and *sex-1*::H1-wCherry transgenic animals. A. Principal component analysis (PCA) plot of lipidomic data showing that *sex-1(y263)* mutants, wild-type (N2) and *sex-1*::H1-wCherry transgenic animals are grouped in different clusters. B. Boxplots showing changes in all lipid classes that are part of lipolysis: triglycerides (TG), diglycerides (DG) and monoglycerides (MG) (n=6 technical replicates per group, 3000 worms per replicate, *P < 0.05, **P < 0.01, ***P < 0.001, ****P < 0.0001; one-way analysis of variance (ANOVA)). C. Boxplots showing changes in levels of lysophosphatidylethanolamine (LPE) and lysophosphatidylcholine (LPC), metabolites involved in phospholipid metabolism, as well as sphingomyelin (SM), derived from sphingolipid metabolism (n=6 technical replicates per group, 3000 worms per replicate, *P < 0.05, **P < 0.01; ****P < 0.0001; one-way analysis of variance (ANOVA)). D. Boxplots showing changes in levels of lipid species involved in glycerophospholipid metabolism: glycerophosphatidic acid (GPA), glycerophosphorylcholine (GPC), glycerophosphoethanolamine (GPE), glycerophosphoglycerol (GPG), glycosylphosphatidylinositol (GPI) and glycerophosphoserine (GPS) (n=6 technical replicates per group, 3000 worms per replicate, *P < 0.05, ***P < 0.001; ****P < 0.0001; one-way analysis of variance (ANOVA)). E. Expression analysis of *sex-1* by RT qPCR in wild-type (N2) animals treated with control or *sex-1* RNAi (n=3 independent experiments, *P < 0.05; two-tailed unpaired t-test). F. Principle component analysis (PCA) plot of lipidomic analysis showing that wild-type (N2) animals treated with control or *sex-1* RNAi are grouped in different clusters. G. Heatmap of lipidomic analysis in wild-type (N2) animals treated with control or *sex-1* RNAi showing differential abundance across all lipid classes. Colour intensity represents the relative changes in lipid levels, with orange indicating upregulation and blue indicating downregulation. H. Quantification of total triglyceride (TG) levels normalised to total protein levels in wild-type (N2) animals treated with control or *sex-1* RNAi (n=3 independent experiments, *P < 0.05; two-tailed unpaired t-test). I. Representative images of *sex-1(y263)* mutants, wild-type (N2) and *sex-1*::H1-wCherry transgenic animals stained with Oil Red O (Images were acquired using a ×10 objective lens, n=1 experiment). J. Quantification of integrated density in the Oil Red O staining as shown in (I) (n=1 experiment, 30 animals, ****P < 0.0001; Kruskal-Wallis test). K. Representative images of day one of adulthood *sex-1(y263)* mutants and wild-type (N2) animals treated with control or *sex-1* RNAi for two generations and stained with Oil Red O (Images were acquired using a ×10 objective lens, n=1 experiment).

**Supplementary Figure 4.**
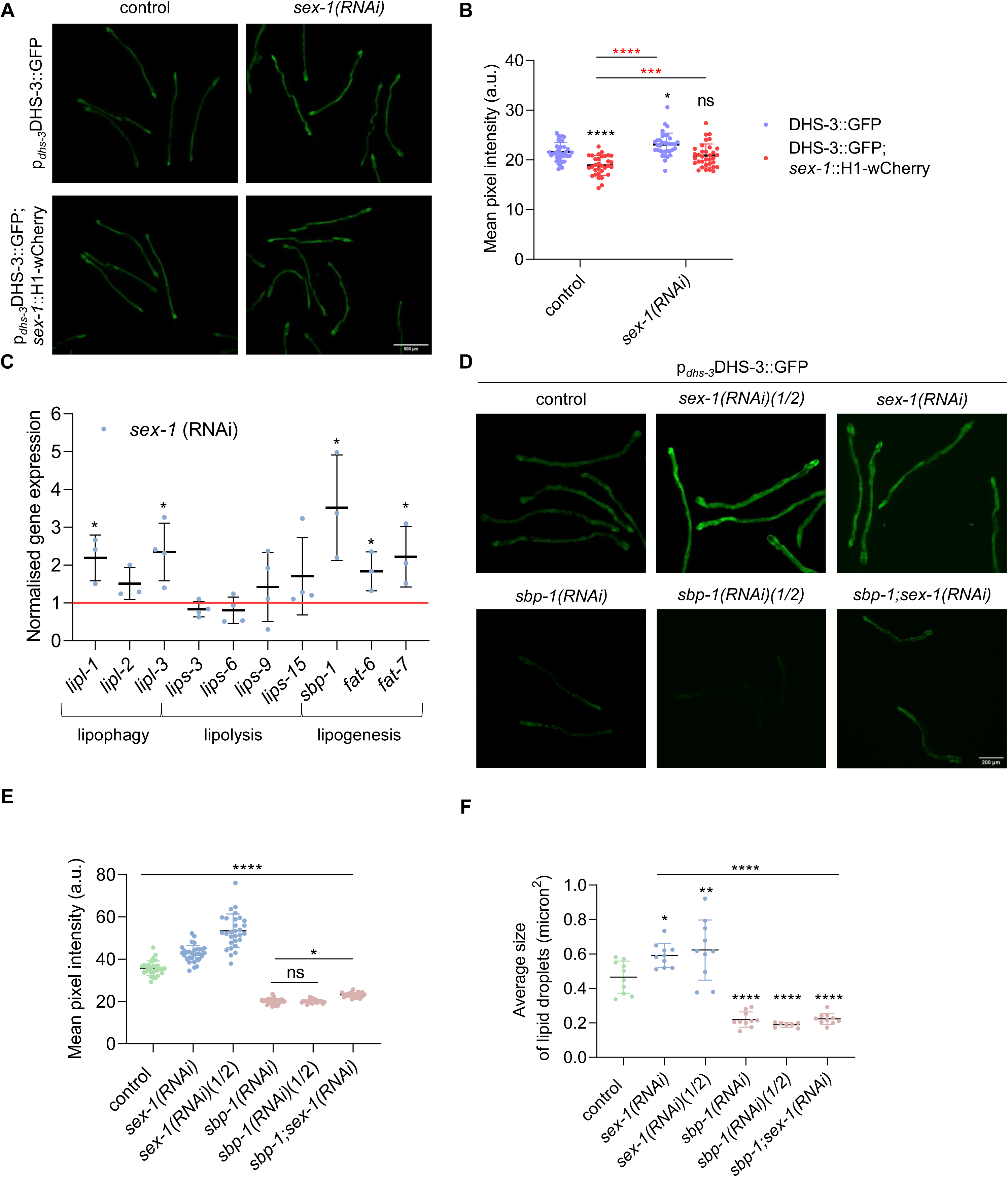
SEX-1 regulates intestinal lipid droplet abundance through SBP-1. A. Representative images of transgenic *sex-1*::H1-wCherry and/or p*_dhs-3_*DHS-3::GFP animals treated with control or *sex-1* RNAi (Images were acquired using a ×4 objective lens, n=2 independent experiments). B. Quantification of the mean GFP signal as shown in (A) (n=2 independent experiments, ***P < 0.001, ****P < 0.0001; one-way analysis of variance (ANOVA)). C. Expression analysis of lipophagy genes: *lipl-1, lipl-2, lipl-3,* lipolysis genes: *lips-3, lips-9, lips-15* and lipogenesis genes: *sbp-1, fat-6, fat-7* by RT qPCR in wild-type (N2) animals treated with control or *sex-1* RNAi (n=3 independent experiments, *P < 0.05; two-tailed unpaired t-test). D. Representative images of transgenic p*_dhs-3_*DHS-3::GFP animals treated with control, *sbp-1*, *sbp-1(1/2)*, *sex-1(1/2)*, *sbp-1;sex-1* or *sex-1* RNAi (Images were acquired using a ×4 objective lens, n=3 independent experiments). E. Quantification of the mean GFP signal as shown in (D) (n=3 independent experiments, *P < 0.05, ****P < 0.0001; one-way analysis of variance (ANOVA)). F. Average size of intestinal lipid droplets assessed by fluorescence as shown in **Figure2.E** following treatment with control, *sbp-1*, *sex-1*, *sex-1(1/2), sbp-1(1/2)* or *sbp-1;sex-1* RNAi (n=1 experiment, 10 animals, *P < 0.05, ****P < 0.0001; one-way analysis of variance (ANOVA)). Each dot represents the average size of lipid droplets measured in the intestine of an individual worm.

**Supplementary Figure 5.**
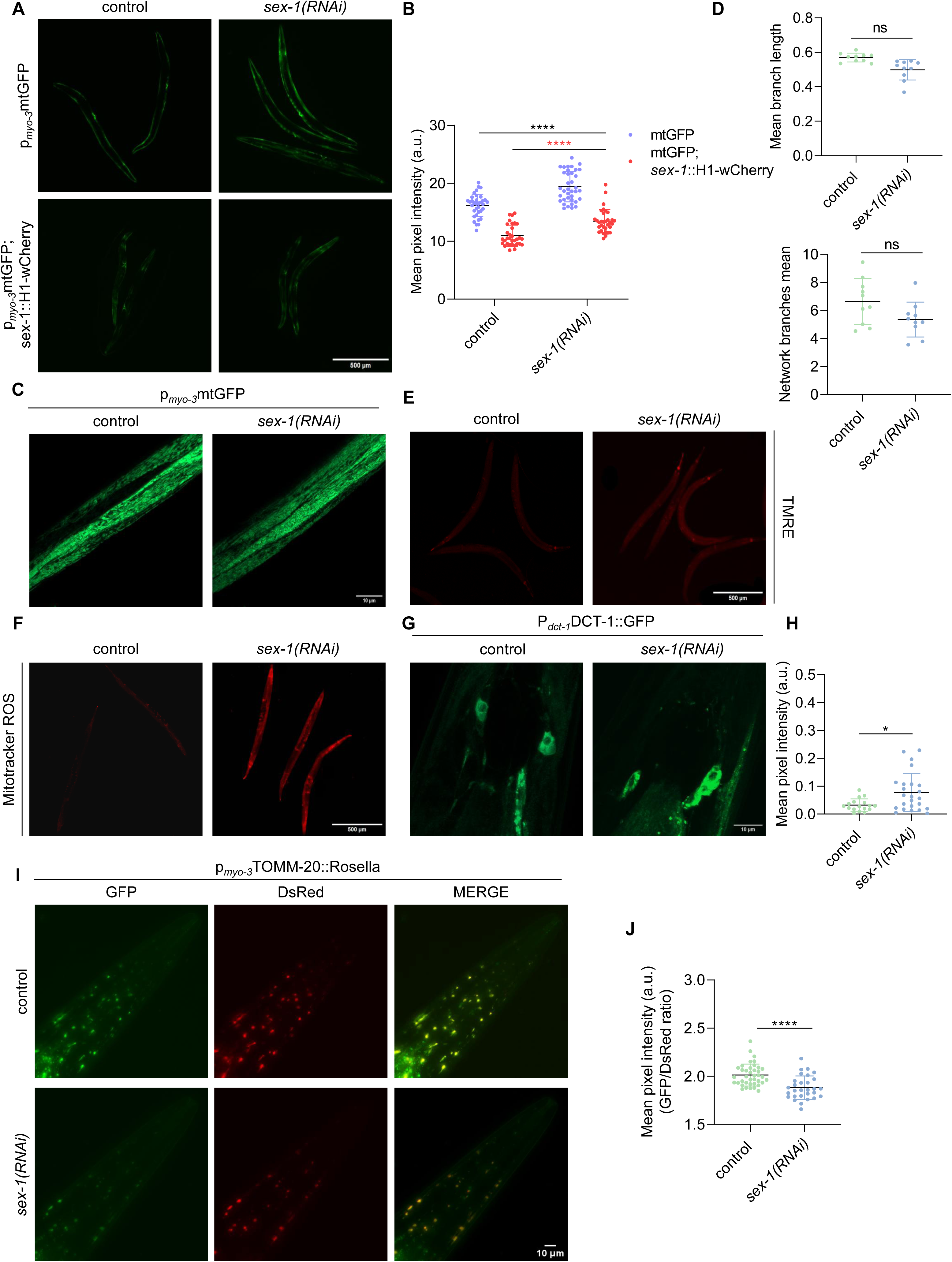
SEX-1 regulates mitochondrial abundance and mitophagy in body wall muscles. A. Representative images of transgenic *sex-1*::H1-wCherry and/or p*_myo-3_*mtGFP animals treated with control or *sex-1* RNAi showing mitochondrial mass in the body wall muscle (Images were acquired using an ×4 objective lens, n=3 independent experiments). B. Quantification of the mean GFP pixel signal in the body wall muscle mitochondrial matrix of animals as shown in (A) (n=3 independent experiments, ***P < 0.001; one-way analysis of variance (ANOVA)). C. Representative images showing mitochondrial morphology in transgenic animals that express mitochondria-targeted GFP in the body wall muscle treated with control or *sex-1* RNAi (Images were acquired using a ×63 objective lens, n=3 independent experiments). D. Quantification of the mean branch length and network branches in transgenic p*_myo-3_*mtGFP animals as shown in (C) (n=3 independent experiments, two-tailed unpaired t-test). E. Representative images of TMRE staining in wild-type (N2) animals treated with control or *sex-1* RNAi, indicating changed in mitochondrial membrane potential (Images were acquired using a ×4 objective lens, n=3 independent experiments). F. Representative images of Mitotracker ROS staining in wild-type (N2) animals treated with control or *sex-1* RNAi showing mitochondrial ROS (Images were acquired using a ×4 objective lens, n=3 independent experiments). G. Representative images of transgenic P*_dct-1_*DCT-1::GFP animals treated with control or *sex-1* RNAi showing localisation of DCT-1 in neurons (Images were acquired using a ×63 objective lens, n=3 independent experiments) H. Quantification of the mean GFP pixel signal of DCT-1 in the head of transgenic P*_dct-1_*DCT-1::GFP animals (n=3 independent experiments, *P < 0.05; Mann-Whitney test). I. Representative images of transgenic animals expressing mitochondria-targeted Rosella (mtRosella) biosensor in the body wall muscle treated with control or *sex-1* RNAi (Images were acquired using a ×20 objective lens, n=3 independent experiments). J. Quantification of the ratio of pH-sensitive GFP to pH-insensitive DsRed in control and *sex-1* RNAi-treated animals. Mitophagy stimulation is indicated by a decrease in the GFP/DsRed ratio (n=3 independent experiments, ****P < 0.0001; Mann-Whitney unpaired test).

**Supplementary Figure 6.**
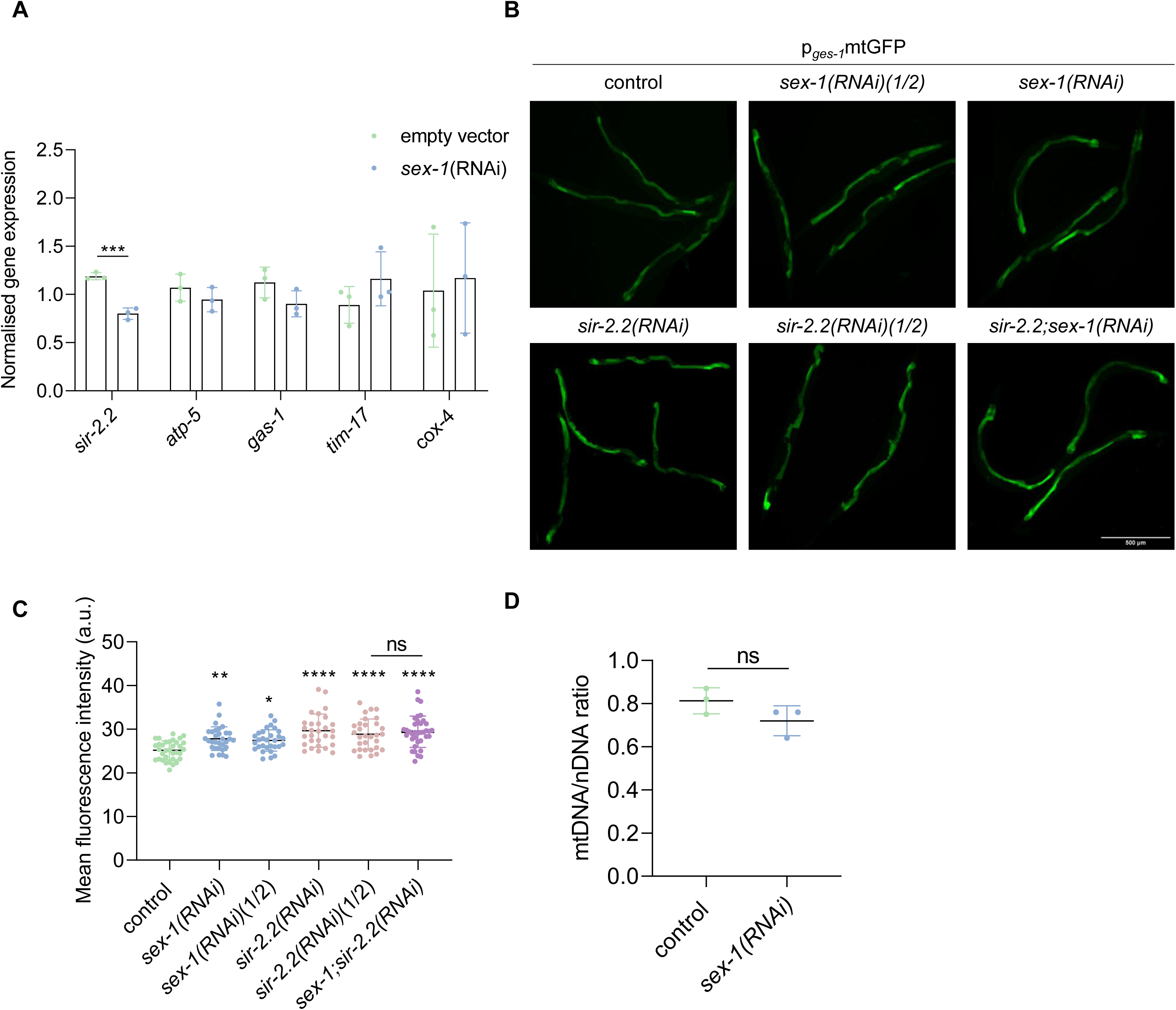
SEX-1 modulates mitochondrial abundance through *sir-2.2.* A. Expression analysis of key mitochondrial quality control genes: *sir-2.2*, *atp-5*, *gas-1*, *tim-17*, *cox-4* by RT qPCR in wild-type (N2) animals treated with control or *sex-1* RNAi (n=3 independent experiments, ***P < 0.001; two-tailed unpaired t-test). B. Representative images of transgenic p*_ges-1_*mtGFP animals treated with control*, sir-2.2*, *sir-2.2(1/2)*, *sex-1(1/2)*, *sir-2.2;sex-1* or *sex-1* RNAi (Images were acquired using a ×4 objective lens, n=3 independent experiments). C. Quantification of the mean GFP pixel signal in the intestinal mitochondrial matrix of animals following treatment with control, *sir-2.2*, *sex-1*, *sex-1(1/2)*, *sir-2.2(1/2)* or *sir-2.2;sex-1* RNAi (n=3 independent experiments, 90 animals in total, ***P < 0.001; Kruskal-Wallis test). D. Quantification of the mtDNA/nDNA ratio under control conditions and upon genetic inhibition of *sex-1* (n=3 independent experiments; two-tailed unpaired t-test).

**Supplementary Figure 7.**
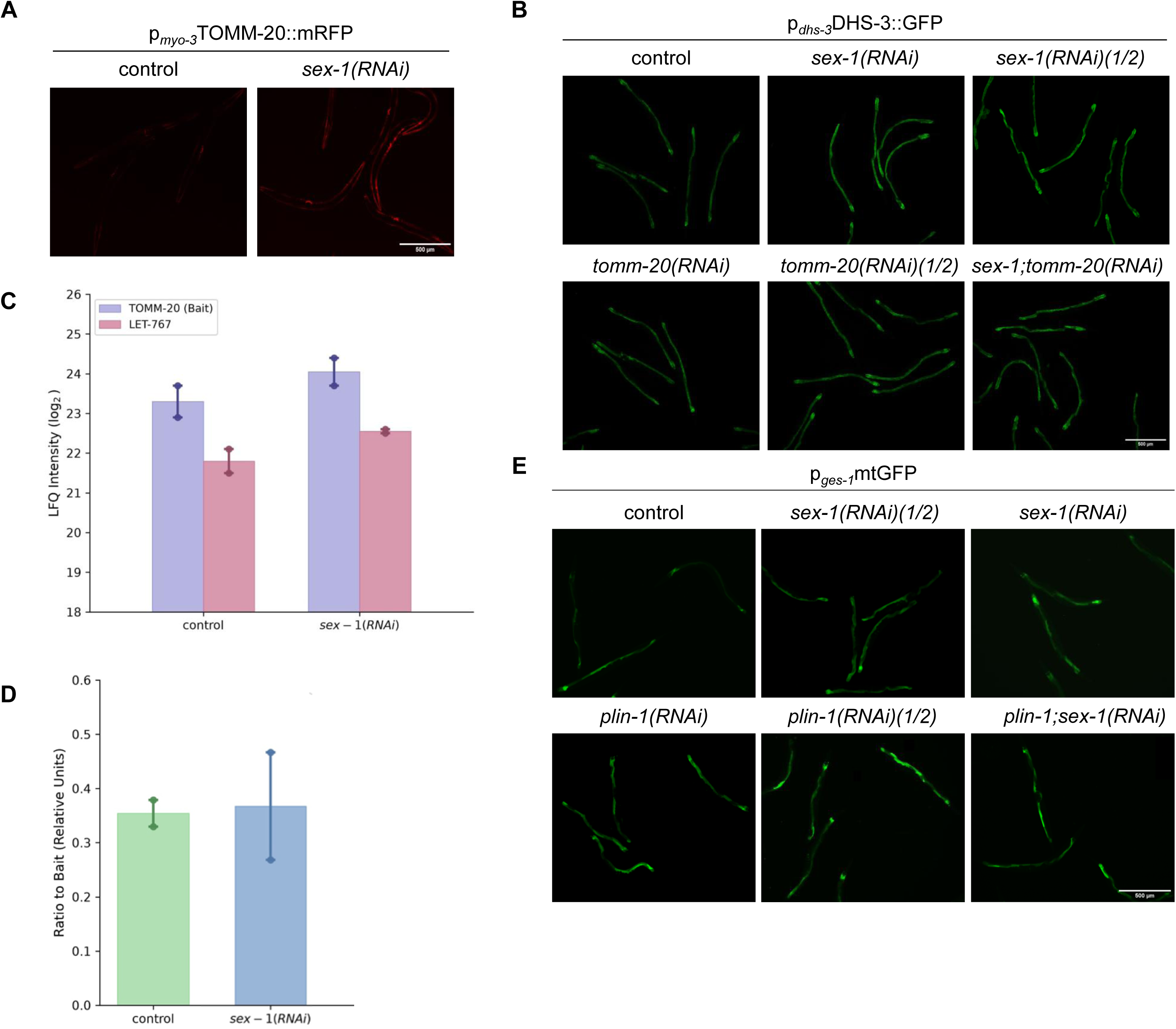
Mediators of mitochondria-lipid droplet interactions identified following SEX-1 depletion. A. Representative images of transgenic p*_myo-3_*TOMM-20::mRFP animals expressing the outer mitochondrial membrane (OMM) protein TOMM-20 treated with control or *sex-1* RNAi (Images were acquired using a ×4 objective lens, n=3 independent experiments). B. Representative images of transgenic p*_dhs-3_*DHS-3::GFP animals treated with control*, tomm-20*, *tomm-20(1/2)*, *sex-1(1/2)*, *tomm-20;sex-1* or *sex-1* RNAi (Images were acquired using a ×4 objective lens, n=3 independent experiments). C. Raw label-free quantification (LFQ) abundance (log_2_ intensity) of bait TOMM-20 and the LET-767 coprecipitated from transgenic p*_myo-3_*TOMM-20::mRFP animals treated with control or *sex-1* RNAi (n = 2 independent biological replicates represented by individual data points). D. Stoichiometric ratio of LET-767 normalised to TOMM-20 bait recovery (LFQ _LET-767_/LFQ _TOMM-20_) in control and *sex-1* RNAi conditions. E. Representative images of transgenic p*_ges-1_*mtGFP animals treated with control*, plin-1*, *plin-1(1/2)*, *sex-1(1/2)*, *plin-1;sex-1* or *sex-1* RNAi (Images were acquired using a ×4 objective lens, n=3 independent experiments).

**Supplementary Figure 8.**
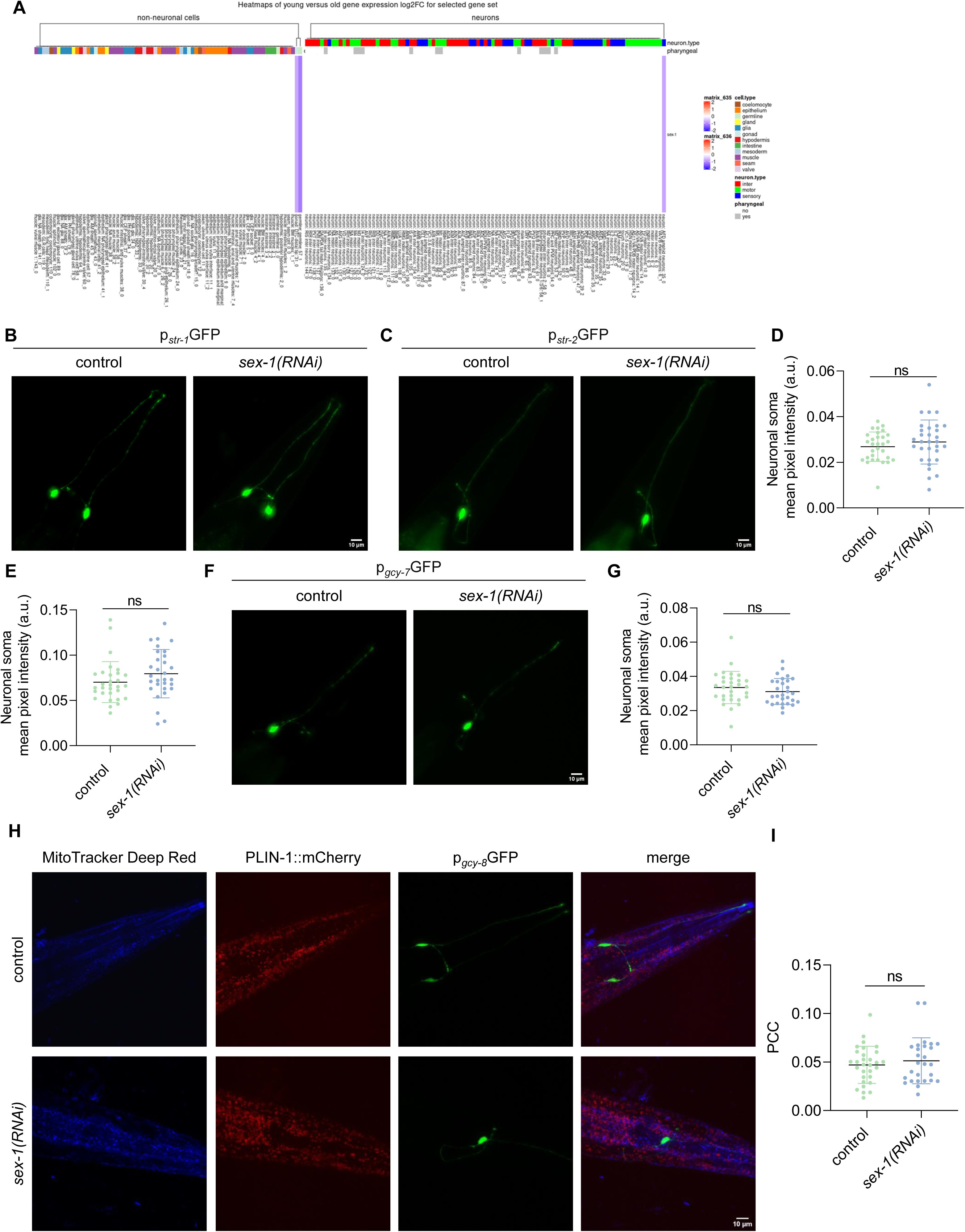
SEX-1 function in amphid neurons. A. Heatmap of the non-neuronal and neuronal cells in which *sex-1* is downregulated with age according to the website: the complete cell atlas of *C. elegans* aging. B. Representative images of transgenic p*_str-1_*GFP animals treated with control or *sex-1* RNAi showing the morphology of AWB neurons (Images were acquired using a ×4 objective lens, n=1 experiment). C. Quantification of the mean GFP signal in AWB neuronal somata as shown in (F) (n=1 experiment, 30 animals, two-tailed unpaired t-test). D. Representative images of transgenic p*_str-2_*GFP animals treated with control or *sex-1* RNAi showing the morphology of AWC neurons (Images were acquired using a ×4 objective lens, n=1 experiment). E. Quantification of the mean GFP signal in AWC neuronal somata as shown in (H) (n=1 experiment, 30 animals, two-tailed unpaired t-test). F. Representative images of transgenic p*_gcy-7_*GFP animals treated with control or *sex-1* RNAi showing the morphology of ASEL neurons (Images were acquired using a ×4 objective lens, n=1 experiment). G. Quantification of the mean GFP signal in ASEL neuronal somata as shown in (F) (n=1 experiment, 30 animals, two-tailed unpaired t-test). H. Representative images of transgenic p*_gcy-8_*GFP animals expressing PLIN-1 (mCherry) treated with control or *sex-1* RNAi and MitoTracker Deep Red (Images were acquired using a ×63 objective lens, n=3 independent experiments) I. Pearson’s correlation coefficient values representing the degree of colocalization between the AFD neuronal reporter and Mitotracker-stained mitochondria (n=3 independent experiments, 30 animals in total)

**Supplementary Figure 9.**
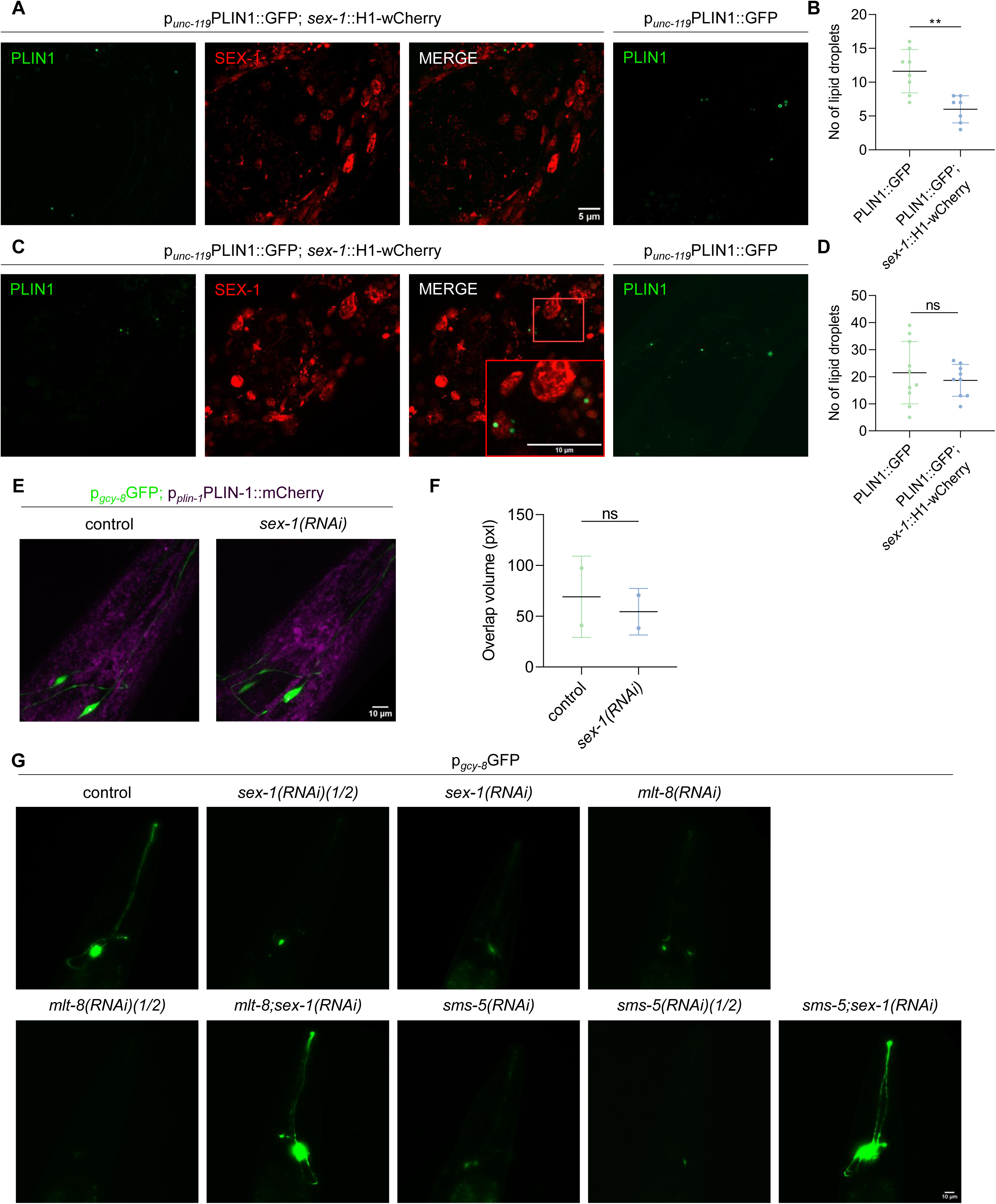
SEX-1 modulates neuronal lipid droplet dynamics. A. Representative images of transgenic *sex-1*::H1-wCherry and/or p*_unc-119_*PLIN1::GFP animals showing lipid droplets in the nerve ring at day 1 of adulthood (Images were acquired using a ×63 objective lens, n=3 independent experiments). B. Number of lipid droplets in the nerve ring of transgenic animals as shown in (A) (n=3 independent experiments, 30 animals in total, **P < 0.01; two-tailed unpaired t-test). C. Representative images of transgenic *sex-1*::H1-wCherry and/or p*_unc-119_*PLIN1::GFP animals showing lipid droplets in the nerve ring at day 5 of adulthood (Images were acquired using a ×63 objective lens, n=3 independent experiments). D. Number of lipid droplets in the nerve ring of transgenic animals as shown in (C) (n = 3 independent experiments, 30 animals in total; two-tailed unpaired t-test). E. Representative images of transgenic p*_gcy-8_*GFP animals expressing PLIN-1 (mCherry), treated with control or *sex-1* RNAi at day 5 of adulthood (Images were acquired using a ×63 objective lens, n=3 independent experiments). F. Quantification of the overlap between lipid droplets and AFD neurons (n=2 independent experiments). G. Representative images of transgenic p*_gcy-8_*GFP animals treated with control, *sex-1*, *sex-1(1/2)*, *mlt-8*, *mlt-8(1/2)*, *mlt-8;sex-1*, *sms-5*, *sms-5(1/2)* or *sms-5;sex-1* RNAi (Images were acquired using a ×20 objective lens, n=3 independent experiments).

## Supplementary Movie legends

**Supplementary Movie 1. Docking of rosiglitazone to SEX-1**. Movie of SEX-1 (yellow) binding to rosiglitazone as predicted by AutoDock Vina and displayed with ChimeraX.

**Supplementary Movie 2. Docking of rosiglitazone to PPAR**γ. Movie of PPARγ (blue) binding to the rosiglitazone as predicted by AutoDock Vina and displayed with ChimeraX.

**Supplementary Movie 3. 3D model of SEX-1 bound to oleic acid.** Movie of SEX-1 (grey) binding to the natural ligand oleic acid (OLA1, pink) as predicted by Alphafold3 and displayed with ChimeraX. The protein residues that are in contact with the ligand have been highlighted.

## Supplementary Table legends

**Supplementary Table 1.** Results of a forward BLASTp analysis of human PPARγ against the *C. elegans* proteome.

**Supplementary Table 2.** Results of a reverse BLASTp analysis of SEX-1 against the *H. sapiens* proteome.

**Supplementary Table 3.** Differentially expressed genes identified by RNA-seq in *sex-1(y263)* mutants.

**Supplementary Table 4.** Differentially expressed genes identified by RNA seq in transgenic *sex-1*::H1-wCherry animals.

**Supplementary Table 5.** Proteins identified by immunoprecipitation coupled to mass spectrometry (IP-MS) using p*_myo-3_*TOMM-20::mRFP as bait.

**Supplementary Table 6.** *C. elegans* strains used in the study.

| Strain | Genotype | Source |
| --- | --- | --- |
| N2 | wild type | CGC |
| TY2384 | *sex-1(y263)* | CGC |
| RW11599 | *stIs11599* [*sex-1::H1-wCherry* + *unc-119(+)*] | CGC |
| LIU1 | *ldrls1* [*p<sub>dhs-3</sub>DHS-3::GFP* + *unc-76(+)*] | CGC |
| LIU2 | *ldrls2* [*p<sub>plin-1</sub> PLIN-1::mCherry* + *unc-76(+)*] | CGC |
| xdIs109 | [*p<sub>unc-119</sub>PLIN1::GFP*, *rol-6*] | X. Huang lab |
| IR3213 | *stIs11599* [*sex-1::H1-wCherry* + *unc-119(+)*]; [*p<sub>unc-119</sub>PLIN1::GFP*, *rol-6*] | This paper |
| IR3221 | [*p<sub>unc-119</sub>PLIN1::GFP*, *rol-6*]; *sid-1(pk3321)*; *uls69* [*pCFJ90(p<sub>myo-2</sub>mCherry)* + *p<sub>unc-119</sub> SID-1*] | This paper |
| IR3142 | *stIs11599* [*sex-1::H1-wCherry* + *unc-119(+)*]; *ldrls1* [*p<sub>dhs-3</sub>DHS-3::GFP* + *unc-76(+)*] | This paper |
| SJ4103 | *zcIs14* [*p<sub>myo-3</sub>mitoGFP*] | CGC |
| IR3217 | *stIs11599* [*sex-1::H1-wCherry* + *unc-119(+)*]; *zcIs14* [*p<sub>myo-3</sub>mitoGFP*] | This paper |
| SJ4143 | *zcIs17* [*p<sub>ges-1</sub>mitoGFP*] | CGC |
| IR3214 | *stIs11599* [*sex-1::H1-wCherry* + *unc-119(+)*]; *zcIs17* [*p<sub>ges-1</sub>mitoGFP*] | This paper |
| IR3193 | *drIs2* [*p<sub>plin-1</sub> PLIN-1::mCherry* + *unc-76(+)*]; *zcIs14* [*p<sub>myo-3</sub>mitoGFP*] | This paper |
| IR3326 | *drIs2* [*p<sub>plin-1</sub> PLIN-1::mCherry* + *unc-76(+)*]; *zcIs14* [*p<sub>ges-1</sub>mitoGFP*] | This paper |
| PS6187 | *syEx1155* [*p<sub>myo-3</sub>TOMM-20::mRFP::3xMyc* + *unc-119(+)*] | CGC |
| PY1283 | *oyIs17* [*p<sub>gcy-8</sub>::GFP* + *lin-15(+)*] | CGC |
| IR3245 | *stIs11599* [*sex-1::H1-wCherry* + *unc-119(+)*]; *oyIs17* [*p<sub>gcy-8</sub>::GFP* + *lin-15(+)*] | This paper |
| IR3216 | *drIs2* [*p<sub>plin-1</sub> PLIN-1::mCherry* + *unc-76(+)*]; *oyIs17* [*p<sub>gcy-8</sub>::GFP* + *lin-15(+)*] | This paper |
| IR3345 | *drIs2* [*p<sub>plin-1</sub> PLIN-1::mCherry* + *unc-76(+)*]; *unc-119(ed3) III*; *pwlIs50* [*p<sub>imp-1</sub>LMP-1* + *unc-119(+)*] | This paper |
| IR2180 | N2; [*p<sub>rab-3</sub>DCT-1::GFP*; *p<sub>rab-3</sub>LMP-1::DsRed*; *p<sub>myo-2</sub>GFP*] | N.T. lab |
| IR2539 | *unc-119(ed3)*; *Ex*[*p<sub>myo-3</sub>TOMM-20::Rosella*; *unc-119(+)*] | N.T. lab |
| IR3317 | *rde-1(ne219)*; *kbls7* [*nhx-2p::rde-1* + *rol-6(su1006)*]; *oyIs17* [*p<sub>gcy-8</sub>::GFP* + *lin-15(+)*] | This paper |
| IR1431 | N2;*Ex001*[*p<sub>dct-1</sub>DCT-1::GFP*] | N.T. lab |
| CX3553 | *kyls104* [*p<sub>str-1</sub>::GFP*] | CGC |
| CX3695 | *kyls140* [*p<sub>str-2</sub>::GFP* + *lin-15(+)*] | CGC |
| OH3191 | *otIs3* [*p<sub>gcy-7</sub>::GFP* + *lin-15(+)*] | CGC |
| HT1593 | *unc-119(ed3) III* | CGC |

**Supplementary Table 7.**
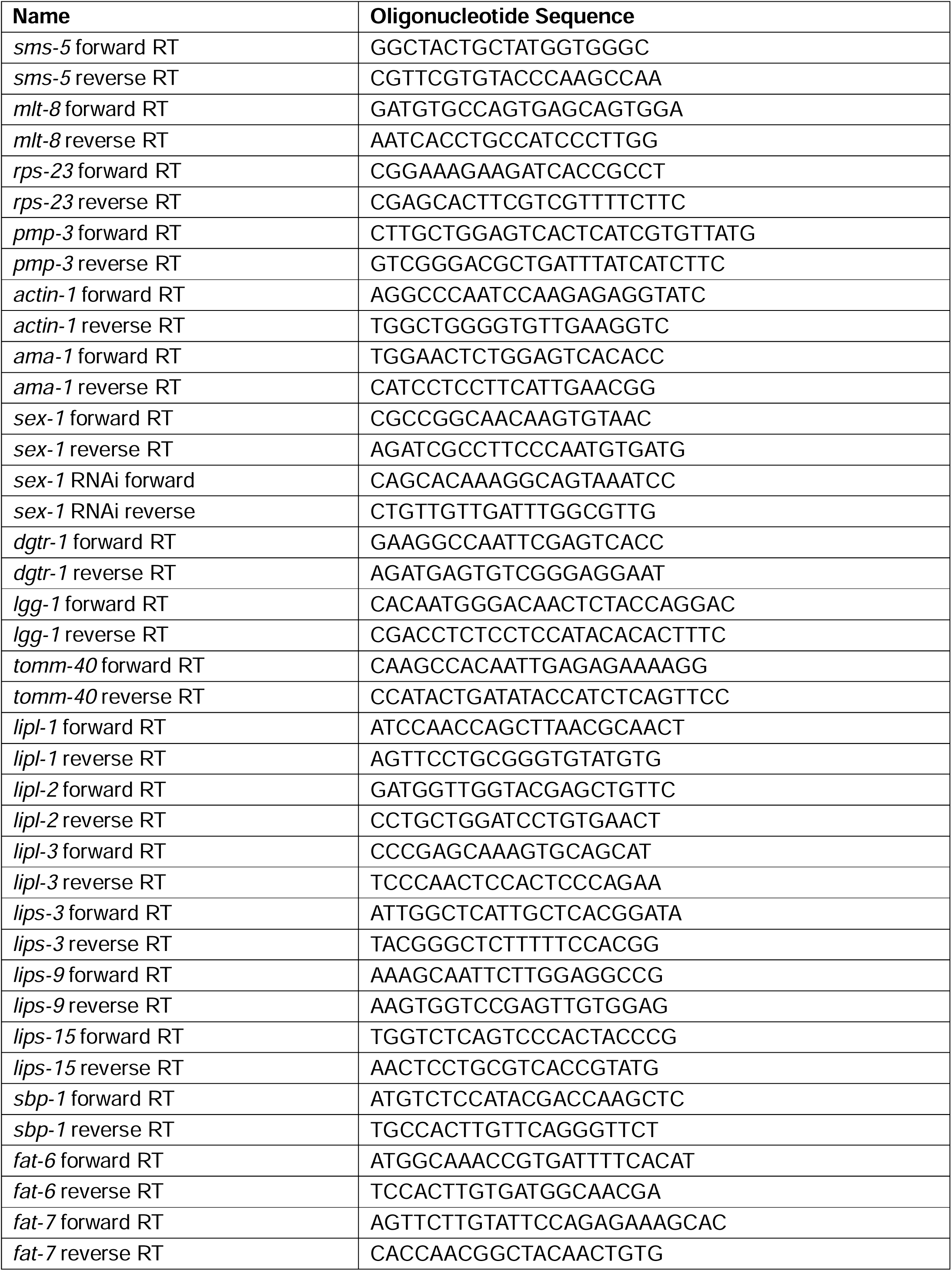
Oligonucleotide sequences used in the study.

**Supplementary Table 8.** Reagents used in the study.

| Reagent | Source | Identifier |
| --- | --- | --- |
| TMRE | Molecular Probes, Invitrogen | T669 |
| Mitotracker Red CM-H2XRos | Molecular Probes, Invitrogen | M7513 |
| Mitotracker Deep Red | Molecular Probes, Invitrogen | M22426 |
| Levamisole | Sigma-Aldrich | 196142 |
| DMSO | AppliChem IWM Reagents | A3672 |
| Roziglitazone | Sigma-Aldrich | R2408 |
| Ampicillin | AppliChem IWM Reagents | A0839 |
| Tetracycline Hydrochloride | AppliChem IWM Reagents | A2228 |
| Oil Red O | Sigma-Aldrich | O0625 |
| RFP antibody | SICGEN | AB1140 |
| PureProteome Protein A/G Mix<br>Magnetic Beads | Sigma-Aldrich/ Merck | LSKMAGAG |

